# Glucocorticoids unleash immune-dependent melanoma control through inhibition of the GARP/TGF-β axis

**DOI:** 10.1101/2025.07.24.666619

**Authors:** Charles H. Earnshaw, Poppy Dunn, Shih-Chieh Chiang, Agrin Moeini, Maria A. Koufaki, Eduardo Bonavita, Massimo Russo, Laetitia Nebot-Bral, Kimberley Hockenhull, Erin Richardson, Anna Pidoux, Charlotte R. Bell, Alexander Baker, Richard Reeves, Robert Sellers, Sudhakar Sahoo, Victoria Fife, Matthew G. Roberts, Theophile Bigirumurame, Caroline Dive, Julia Newton-Bishop, Jérémie Nsengimana, Christopher E. M. Griffiths, Santiago Zelenay

**Author notes:** Department of Biomedical Sciences, Humanitas University; Milan, Italy (Eduardo Bonavita).

## Abstract

Half of advanced melanoma patients fail to benefit from immune checkpoint blockade and novel treatments are urgently required. Testing topical medications used in other skin diseases for anti-cancer activity in an immunotherapy-resistant murine melanoma model, we counterintuitively found that glucocorticoids elicit rapid cytotoxic T lymphocyte (CTL)-dependent tumor control. Genetic ablation of the glucocorticoid receptor in different cellular compartments revealed glucocorticoids acted not on immune cells but directly on tumor cells to downregulate expression of GARP (glycoprotein A repetitions predominant). This inhibited TGF-β signaling and unleashed CTL killing. In agreement, glucocorticoids stimulated tumor control in multiple cancer models, but only if the tumors also responded to pharmacological inhibition of TGF-β signaling. Furthermore, melanoma patients with high glucocorticoid receptor expression or signaling showed improved prognosis and lower TGF-β signaling in tumor-infiltrating CTLs. Additionally, elevated GARP expression correlated with reduced survival, including in immunotherapy-treated patients. Thus, the GARP/TGF-β axis emerges as a glucocorticoid-sensitive cancer cell-intrinsic immune evasive mechanism.

**Significance:** Screening widely used topical treatments in a melanoma model, this study uncovers a surprising role for glucocorticoids in triggering CD8^+^ T cell-dependent tumor control through downregulation of GARP and thus TGF-β signaling. Melanoma patient sample analysis supported these findings suggesting GARP/TGF-β activity functions as a tumor cell-intrinsic immune evasive mechanism, and GARP expression may serve as both a biomarker of poor antitumor immunity and a therapeutic target to improve the response to immunotherapy.

## Introduction

Immune checkpoint blockade (ICB) is the most promising novel pan-cancer treatment approach since development of the first chemotherapies. Unprecedented results have been observed in multiple cancer types including malignancies once thought untreatable [1]. Particularly in melanoma, ICB has transformed treatment outcomes with approximately half of advanced melanoma patients experiencing lasting benefit from the combination of anti-cytotoxic T-lymphocyte-associated protein 4 (anti-CTLA4) and anti-programmed cell death protein 1 (anti-PD-1) therapy [2]. An improved mechanistic understanding of the underlying basis for treatment failure is needed to develop new treatments for the other half of melanoma patients who fail to respond or benefit only short-term.

Prior work has investigated why some patients benefit from immunotherapy, and others do not. Factors such as high tumor mutational burden, high tumor expression of Programmed Death-Ligand 1 (PD-L1), elevated intratumoral CTL infiltration or interferon (IFN) signaling and the presence of tertiary lymphoid structures have been, among many others, associated with an increased likelihood of response to ICB [3–6]. Indeed, given the complex and often disparate inflammatory profiles active in the progression of tumors, a diverse array of cellular and non-cellular immune mediators within the tumor microenvironment have been shown to play critical roles in shaping the response to immunotherapy [7–9]. Response heterogeneity following ICB treatment is seen, just as in patients, across syngeneic mouse cancer models, making them tractable and versatile experimental systems to further identify resistance mechanisms and novel therapeutic targets.

Recent proof-of-principle work has shown that ICB efficacy can be improved through the blockade of other immune checkpoints [10–12] or the addition of chemotherapeutic [13], and immune-stimulatory [14, 15] drugs. Among the latter, we and others have previously shown that anti-inflammatory drugs can boost the immune response to cancer [16–20]. This is consistent with the notion that cancer-associated inflammation, a complex and dynamic process denoting the functional consequences of both innate and adaptive immune responses, can have antagonistic tumor-promoting or -inhibitory effects [7, 8, 21, 22]. Under this premise, we therefore sought to test whether existing widely available and inexpensive topical medications used to treat a variety of dermatoses had anti-tumor activity in ICB-refractory melanoma. These included the drugs 5-fluorouracil (as a chemotherapeutic), imiquimod (as an immune-stimulatory drug), and diclofenac and glucocorticoids (GCs) (as anti-inflammatory drugs; with the latter recently shown to lead to improved ICB-responses in certain preclinical models [17]).

Among the different topical drugs tested, we found that GCs uniquely trigger CD8^+^ T cell-dependent tumor growth control. This was unexpected due to the well-known immunosuppressive functions of GCs. The cancer-inhibitory effects of topical GCs were observed in numerous, but not all, melanoma and non-melanoma cancer models. Mechanistically, we demonstrate that GCs act directly on melanoma cells via the GC receptor (GR) to inhibit expression of GARP (glycoprotein-A repetitions predominant), impairing in turn the activation of the immunosuppressive cytokine transforming growth factor beta (TGF-β). Consistently, GCs only benefited tumor models that also showed sensitivity to inhibition of TGFβ signaling. In agreement with these findings, we show that in melanoma patients, high GR expression or GC signaling at the tumor site associates with improved survival particularly in tumors with elevated CD8^+^ T cell-infiltration. Moreover, we found that GARP is heterogeneously expressed by cancer cells in human melanoma specimens, and that its expression levels negatively correlate with patient survival, outcome from immunotherapy and the activation state of tumor-infiltrating CTLs. Lastly, patients with elevated melanoma cell-intrinsic GR expression have reduced levels of TGF-β signaling in infiltrating CD8^+^ T cells. Together, our findings suggest that GC use has the potential to benefit a subset of immunotherapy-resistant patients, and that direct inhibition of the GARP-TGF-β axis on tumor cells represents a promising new therapeutic target.

## Results

### Topical glucocorticoid treatment stimulates acute tumor control in an immunotherapy-resistant melanoma model

To assess the anti-tumor activity of widely used cream-based medications to treat ICB-resistant melanomas, we first identified a murine syngeneic melanoma model (20967) that grew progressively upon orthotopic intradermal implantation and did not respond to single or dual combination of anti-PD-1 and anti-CTLA4 therapy in wild-type C57BL/6 mice (Fig. 1A, Supplementary Fig. S1A-E). This contrasted with 5555 tumors, which, like other commonly studied pre-clinical melanoma models [16, 23–25], responded vigorously to combination treatment (Fig 1A, Supplementary Fig. S1B).

**Figure 1.**
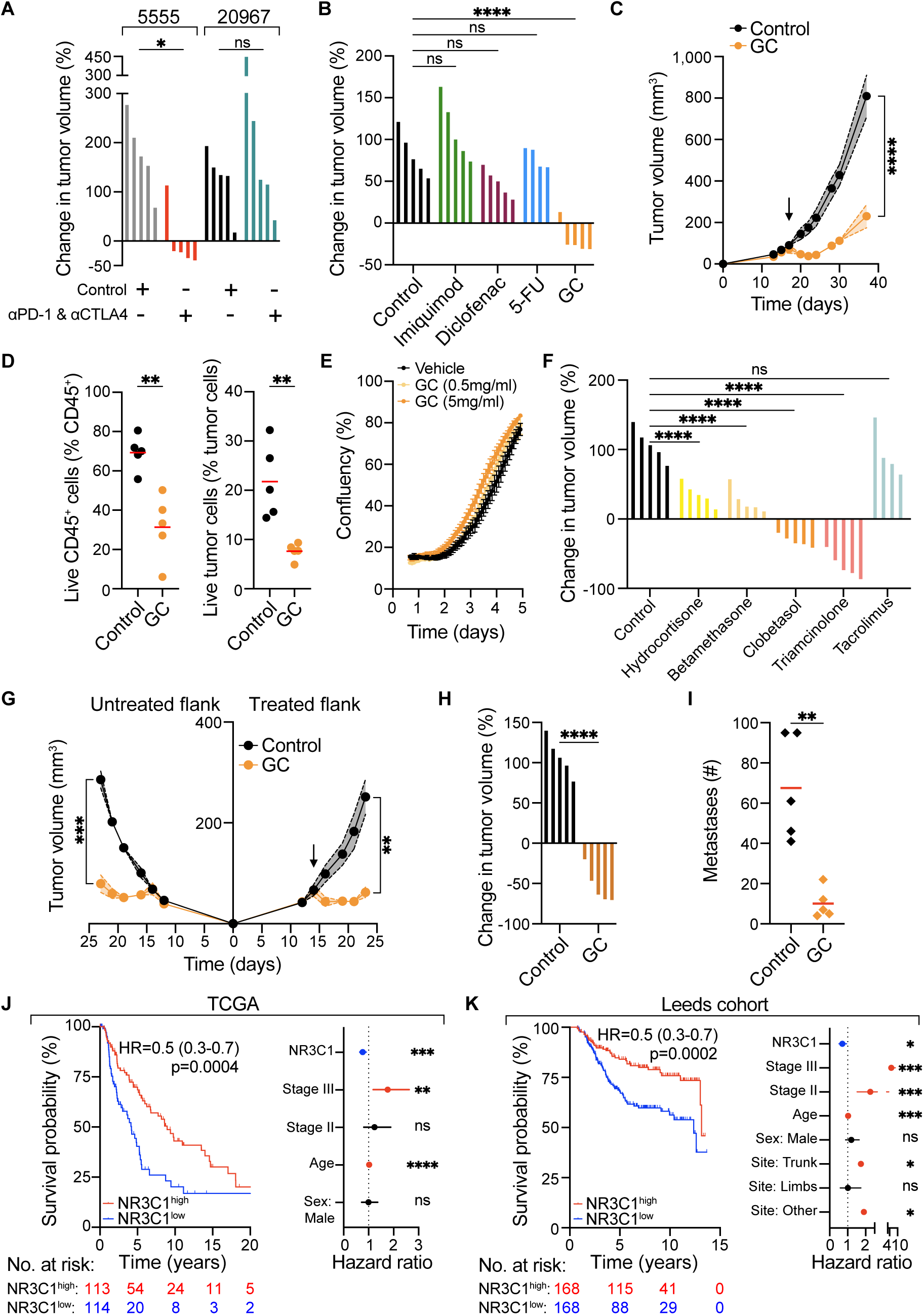
GCs stimulate melanoma control and are associated with good prognosis in patients. (**A**) Waterfall plots showing percentage change in tumor volume at day 5 post-treatment in response to ICB of 5555 and 20967 melanoma tumors (n=5 per group). (**B**) Waterfall plots showing percentage change in tumor volume at day 5 post-treatment of 20967 tumors to different topical drugs (n=4-5 per group). (**C**) Growth profile of 20967 tumors treated with control and GCs (n=5 per group). Arrow indicates start of treatment. (**D**) FACS analysis of GC-treated tumors 5 days after treatment showing live immune cells (left panel) and live tumor cells (right panel) (n=5 per group). (**E**) *In vitro* growth of 20967 melanoma cells treated with different concentrations of GC (clobetasol propionate) measured by IncuCyte. (**F**) Waterfall plot showing percentage change in tumor volume at day 5 post-treatment of 20967 tumors to control, different GCs topically (or, for triamcinolone, intratumourally) or tacrolimus (n=4-5 per group). (**G**) Growth profiles of bilaterally injected melanoma tumors (n=5 per group), with one tumor per mouse treated with control or GC. Arrow indicates start of treatment. (**H**) Waterfall plot showing percentage change in tumor volume at day 5 post-treatment of 20967 tumors with control and GCs administered by intraperitoneal injection (n=5 per group). (**I**) Number of metastases in lungs of mice treated systemically with control or GC (n=5 per group) (**J, K**) Kaplan-Meier survival plots of cutaneous melanoma (TCGA, n=458 and Leeds melanoma cohort, n=703) patients stratified by upper and lower quartile of GR (J, K; left panel), and forest plots showing multivariate COX regression analysis for the indicated factors in TCGA and Leeds melanoma cohort patients (J, K; right panel). Data are expressed as mean ± SEM; one-way ANOVA (A, B, F), two-way ANOVA (C, G), unpaired t-test (D, H) and Mann-Whitney U test (I). Hazard ratio (95% confidence interval), log-rank (Mantel-Cox) test (J, K). *, *P* < 0.05; **, *P* < 0.01; ***, *P* < 0.001; ****, *P* < 0.0001; ns, not significant.

We used this ICB-unresponsive model to screen for tumor-suppressive activity of medications available in topical form and used commonly in the treatment of conditions such as actinic keratosis, squamous and basal cell carcinoma, or inflammatory dermatoses such as eczema and psoriasis. Mice bearing established 20967 tumors received imiquimod, a toll like receptor 7 agonist; 5-fluorouracil (5-FU), a chemotherapy agent; diclofenac, a non-steroidal anti-inflammatory drug; or clobetasol propionate, a potent GC (Supplementary Fig. S2A). Unexpectedly, GCs were the only topical treatment that caused rapid tumor shrinkage and led to profound tumor growth inhibition in all mice (Fig. 1B, C). The acute tumor inhibitory effects of topical GCs were invariably observed in all mice even when treating large established tumors of over 300 mm^3^ (Supplementary Fig. S2B). Analysis of cell viability early on following GC application revealed a significant drop in the fraction of live cells in both melanoma and tumor-infiltrating immune cell compartments indicating that GCs had tumoricidal activity and were not merely locally reducing inflammatory swelling (Fig. 1D). Alongside tumor growth inhibition, topical GCs induced a decrease in blood leukocyte counts and spleen size (Supplementary Fig. S2C). Importantly, clobetasol propionate did not impact the growth of 20967 melanoma cells *in vitro,* arguing against a direct cytotoxic effect of GCs on tumor cells (Fig. 1E).

A range of different GCs showed rapid tumor-suppressive effects *in vivo*, with topical clobetasol propionate and triamcinolone delivered intratumorally eliciting particularly potent reductions in tumor volume of up to 85% following just five days of treatment (Fig. 1F). Tacrolimus, in contrast (an immunosuppressive medication also used in the treatment of skin disease but with a different mechanism of action to GCs) did not alter tumor growth, suggesting the tumor inhibitory effect was GC specific (Fig. 1F).

To assess whether topical GC treatment of tumors elicited a systemic anti-tumor response, we implanted tumors bilaterally and treated one tumor only. This treatment provoked similar tumor growth inhibition in both the treated and untreated contralateral tumor (Fig. 1G). Given this finding and the abrupt reduction in leukocyte counts in blood and spleen (Supplementary Fig. S2C) suggesting GCs were systemically absorbed following topical application, we next sought to evaluate whether systemic administration of GCs would also stimulate tumor control. Indeed, intraperitoneal injection of GCs resulted in similar tumor shrinkage as when GCs were administered topically or intratumorally (Fig. 1H), potentially explaining the apparent abscopal effect observed above. To determine whether the tumor-suppressive effects of GCs were specific to the skin environment, we intravenously injected melanoma cells to model lung metastases and administered GCs systemically. Notably, this treatment led to a significant decrease in metastatic burden (Fig. 1I). Taken together, these findings reveal that GCs of varying potencies and administered via different routes can induce rapid tumor regression both locally and at distant sites.

### Glucocorticoid receptor expression and activity are positively prognostic in human melanoma

To assess the impact of the GC axis in human melanoma, we interrogated The Cancer Genome Atlas database (TCGA). Elevated transcript levels of *NR3C1*, the gene encoding for the GR, were associated with significantly improved overall survival (OS) (median survival 9 years vs 4.2 years, hazard ratio 0.5, p=0.0004, Fig. 1J). The beneficial effect of elevated GR expression in melanoma was recapitulated in an independent melanoma patient cohort [26] (Fig. 1K). This cohort comprises primary cutaneous melanoma specimens as opposed to the TCGA melanoma dataset, which consists mainly of metastatic samples, explaining the OS difference. Importantly, the prognostic utility of GR expression was independent of stage, gender and age in both patient datasets (Fig. 1J, K). *NR3C1* mRNA levels positively correlated with a GC-stimulated gene signature (described later: Supplementary Tables S1 and S2) arguing that higher tumor GR expression also means increased intratumoral GC activity (Supplementary Fig. S3A). Elevated levels of the GC-stimulated gene signature, as well as an independently derived signature of GC activity [27], were also associated with improved OS in melanoma (Supplementary Fig. S3B, C). We additionally examined the GC synthesis pathway whereby the enzyme 11β-HSD1 drives production of active GCs and 11β-HSD2 inactivates GCs (Supplementary Fig. S3D). In keeping with a beneficial effect of intratumoral GCs in melanoma, transcript levels of the genes encoding for 11β-HSD1 and 11β-HSD2 were positively and negatively associated, respectively, with patient survival (Supplementary Fig. S3E, F). Overall, and in agreement with our mouse findings, this analysis suggests that in human melanoma, increased GC signaling via the GC receptor associates with improved survival.

### GC-induced tumor control is CD8^+^ T cell dependent

To investigate the mechanism underlying the acute tumor control induced by topical GCs, we first queried whether the effect of GC treatment relied on the immune status of the host. Remarkably, tumor growth inhibition following GC administration was lost in *Rag1^-/-^*mice, lacking T and B cells, or in wild-type mice depleted of CD8^+^ T cells, but not CD4^+^ T cells (Fig. 2A-D), demonstrating that adaptive immunity, and specifically CTLs, were key mediators of the anti-tumor effect of GCs.

**Figure 2.**
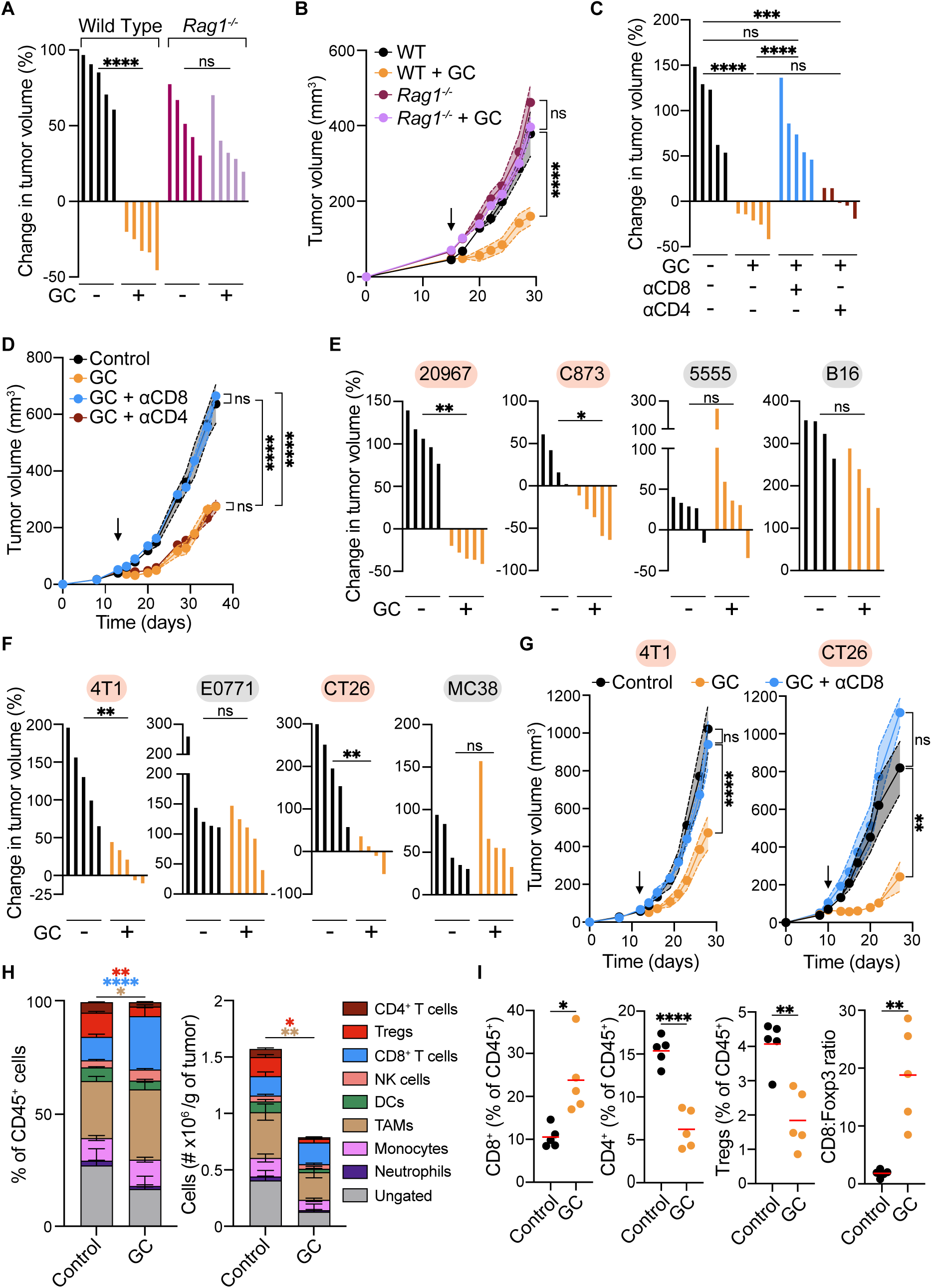
The tumor-suppressive effect of GCs is CD8^+^ T cell-dependent and seen in select tumor models. (**A, B**) Waterfall plot showing percentage change in tumor volume at day 5 post-treatment (A), and full growth profile (B) in wild type or Rag1^-/-^ mice treated with control or GCs (n=5 per group). (**C, D**) Waterfall plots showing percentage change in tumor volume at day 5 post-treatment (C), and full growth profile (D) in mice depleted of CD4^+^ or CD8^+^ cells and treated with GCs (n=5 per group). (**E, F**) Waterfall plots showing percentage change in tumor volume at day 5 post-treatment of 20967, 5555, C873 and B16 melanomas (E) and 4T1, E0771, CT26 and MC38 tumors (F) to GCs (n=4-5 per group). Red highlight indicates GC-responsive tumors and grey highlight indicates GC-unresponsive tumors. (**G**) Growth profile of 4T1 tumors (left) or CT26 (right) implanted into control mice or mice depleted of CD8^+^ cells treated with control or GCs (n=5 per group). (**H**) Percentage of total leukocytes (left panel) and number per gram of tumor (right panel) in control and GC-treated 20967 melanomas (n=5 per group) (**I**) Intratumoral immune-infiltrate analysis of 20967 melanomas on day 5 post-treatment with control or GCs (n=5 per group). Data are expressed as mean ± SEM; one-way ANOVA (A, C), two-way ANOVA (B, D, G), Mann-Whitney test (E, F) or unpaired t-test (H, I). *, *P* < 0.05; **, *P* < 0.01; ***, *P* < 0.001; ****, *P* < 0.0001; ns, not significant. Arrow indicates start of GC treatment.

To assess whether GCs stimulated tumor control in other melanoma and non-melanoma tumors, we next evaluated diverse additional murine cancer models: three melanoma (5555, C873, and B16), two breast cancer (4T1 and E0771), and two colorectal cancer (CT26 and MC38) models. Tumors formed by C873 melanoma cells responded vigorously to GCs like 20967 tumors, whereas the growth of 5555 and B16 tumors was not affected (Fig. 2E). Of the breast and colorectal cancer models, 4T1 and CT26, but not E0771 or MC38, responded to GC treatment (Fig. 2F) and this effect, as for 20967 tumors, required the presence of CD8^+^ T cells (Fig. 2G). Thus, the CTL-dependent tumor-suppressive effect of topical GCs can be observed in multiple, but not all, cancer models.

### GCs deplete conventional and regulatory CD4^+^ T cells but spare tumor infiltrating CTLs

Next, we examined how the immune infiltrate was altered in GC-treated 20967 tumors. The overall number of both lymphoid and myeloid cells was substantially reduced following treatment, with a particularly pronounced decrease in total CD4^+^ T cells and CD4^+^ Foxp3^+^ regulatory T cells (Tregs) (Fig. 2H, I, Supplementary Fig. S4, and S5A). In sharp contrast, CD8^+^ T cells were spared from the deleterious effects of GC treatment (Fig. 2H, I, Supplementary Fig. S5A-C), a finding previously reported in the skin of psoriasis patients upon treatment with the same GC [28]. This resulted in a two-fold increase in the frequency of CD8^+^ T cells and a ∼9-fold increase in the ratio of CD8^+^ T cells to Tregs (Fig. 2I). Of note, however, antibody-mediated Treg depletion did not noticeably impair tumor growth (Supplementary Fig. S5D, E), suggesting Treg ablation is not sufficient to account for the GC-driven cytotoxic T lymphocyte (CTL)- mediated tumor control.

Further phenotypic characterization of tumor-infiltrating CTLs by flow cytometry indicated that the proportion of stem cell-like PD-1^+^ TCF1^+^ or dysfunctional PD-1^+^ TOX^+^ cells was not altered by GC administration, whereas naïve PD-1^-^ TCF1^+^ CD8^+^ T cells were reduced (Fig. 3A). Still, altogether, these CD8^+^ T cell subsets did not account for more than ∼15% of the total tumor-infiltrating CTLs. Based on their expression of CD69 and CD103, most tumor-infiltrating CD8^+^ T cells resembled tissue-resident memory cells (Supplementary Fig. S5A). Whereas GC treatment did not alter the proportion of these cells, it augmented their expression of CD69 and PD-1 and decreased CD39 levels (Fig. 3B). With respect to cytotoxic effector functions, the frequency of IFNγ or TNF-producing CD8^+^ T cells remained unchanged following GC administration (Supplementary Fig. S5B). However, we observed a strong correlation between the extent of tumor control and the frequency of IFNγ^+^ or TNF^+^ CTLs, specifically in the GC treated group (Supplementary Fig. S5C). Together, these results suggest that although CD8^+^ T cells in GC-treated tumors phenotypically resemble those in control-treated tumors, their relative abundance is significantly increased, and their effector function is closely linked to the degree of tumor regression.

**Figure 3.**
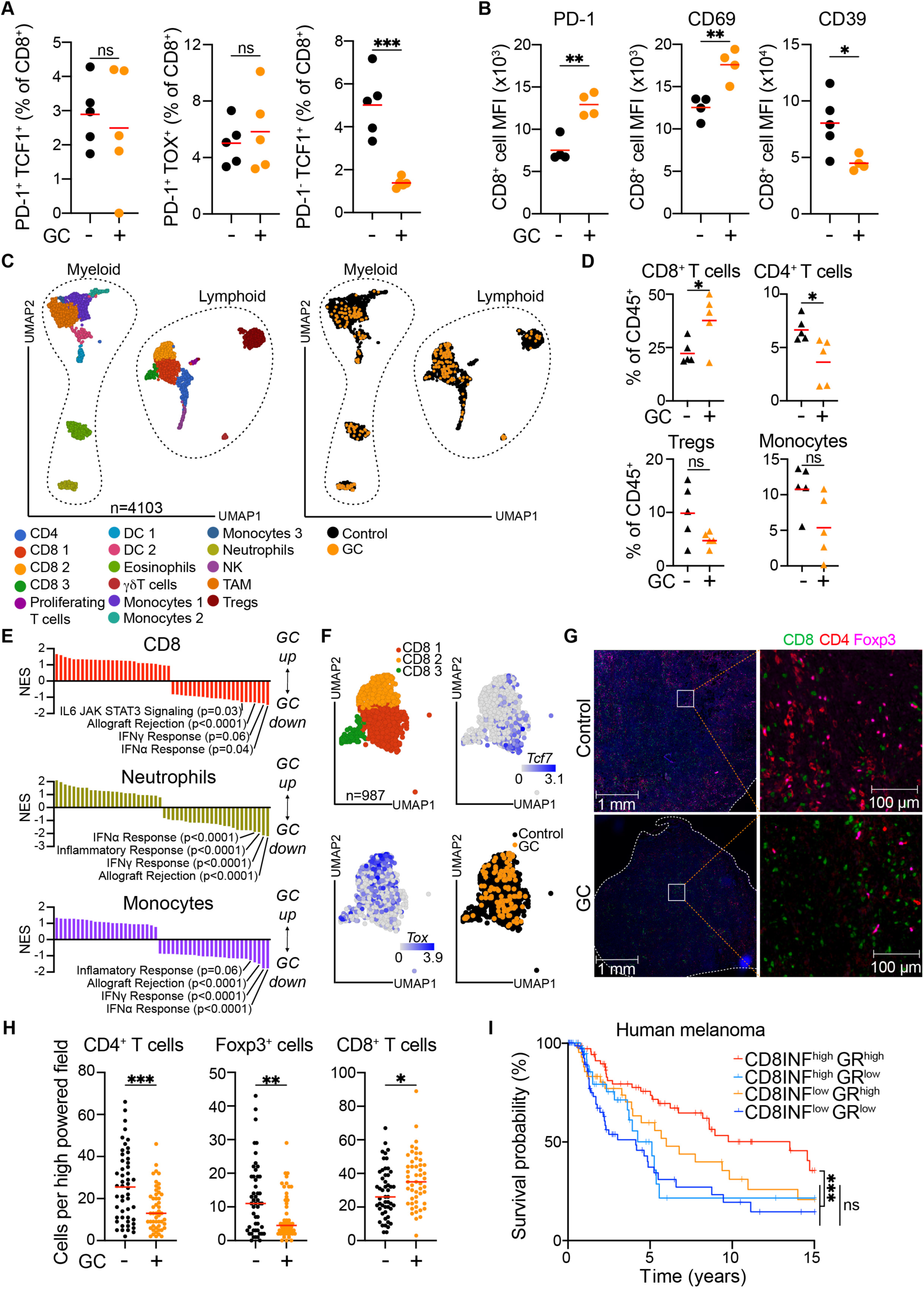
Tumor-infiltrating CD8^+^ T cells are acutely enriched following GC-treatment. (**A, B**) Intratumoral immune-infiltrate analysis of 20967 melanomas on day 5 post-treatment with control or GC (n=4-5 per group). (**C**) Unified Manifold Approximation and Projection (UMAP) of 10 individually hashtagged 20967 melanoma tumors analyzed by single-cell RNA sequencing (scRNA-seq), treated with control or GCs (n=5 per group). Annotations by cell cluster (left panel) and treatment (right panel) shown. (**D**) Immune infiltrate analysis of scRNA-seq performed on 20967 melanoma tumors 4 days post-treatment with GC (n=5 per group). (**E**) GSEA of different immune cell populations in GC-treated tumors, showing Normalized Enrichment Scores (NES). (**F**) UMAPs showing CD8^+^ T cells classified by indicated clusters, gene expression or drug treatment. (**G, H**) Immunofluorescence analysis of control and GC treated tumors. Representative image of multiplex immunofluorescence staining of tumor sections showing CD8^+^, Foxp3^+^ and CD4^+^ T cells (G), and quantification in 10 independent fields (H) from 5 mice, 5 days on treatment. (**I**) Survival analysis of TCGA melanoma patients split by both GR expression and CD8^+^ T cell infiltration (CD8^+^ INF) determined by MCP counter (n=458). Data are expressed as mean; unpaired t-test (A, B, D, H) or log-rank (Mantel-Cox) test with Bonferroni correction (I). *, *P* < 0.05; **, *P* < 0.01; ***, *P* < 0.001; ns, not significant.

Single cell RNA sequencing of tumor-infiltrating CD45^+^ cells from individually hash-tagged tumors further confirmed the above data, showing no major change in the overall myeloid and lymphoid infiltrate composition apart from an increase in CD8^+^ T cells and a reduction in CD4^+^ T cells upon GC treatment (Fig. 3C, D). Gene Set Enrichment Analysis (GSEA) of independent cell clusters showed a consistent decrease in ‘Inflammatory Response’, ‘IFN-γ Response’, ‘IFN-α Response’ and ‘Allograft Rejection’ Hallmark Gene sets in both lymphoid and myeloid cells from GC-treated tumors, consistent with the anticipated broad anti-inflammatory and immunosuppressive effect of GCs (Fig. 3E). Among the three major clusters containing CD8^+^ T cells, clusters 2 and 3 were enriched in more ‘exhausted’ cells with higher *Tox* expression, and cluster 1 in ‘stem-like’ cells with higher *Tcf7* expression (Fig. 3F). GC treatment led to no apparent enrichment or depletion in any of these independent CD8^+^ T cell transcriptional states (Fig. 3F). Examination of tumor-infiltrating T cells by immunofluorescence microscopy confirmed that GC treatment led to an abrupt depletion of CD4^+^ T cells and Tregs but spared CTLs, with no clear morphological tissue alterations (Fig. 3G, H, Supplementary Fig. S6A-C). CTLs were relatively evenly distributed across untreated 20967 tumors, and their density marginally increased following GC treatment, again illustrating a markedly elevated CD8^+^:Treg ratio (Fig. 3G, H, Supplementary Fig. S6C). Taken together, immune infiltrate profiling revealed that GC treatment resulted in rapid enrichment of CD8^+^ T cells, most of which were tissue-resident memory cells, without any obvious alterations in their phenotype or transcriptional state.

We next queried if the favorable prognosis of melanoma patients with higher GR expression was similarly linked to intratumoral CTL presence. To interrogate this, we evaluated the survival of melanoma patients stratified based on both intratumoral GR levels and inferred CD8^+^ T cell infiltration. Notably, patients with both high GR expression and high CD8^+^ T cell infiltration survived more than twice as long as those with high CD8^+^ T cells but low GR expression (Fig. 3I). Indeed, the outcome of the latter group was comparable to that of the two CD8^+^ T cell low patient groups, irrespective of their GR levels. Therefore, these data suggest the association of GR expression with improved prognosis in melanoma patients is intimately linked to having elevated CD8^+^ T cell infiltration and demonstrate, in keeping with the mouse data, a beneficial CTL-dependent contribution of intratumoral GC activity in human melanoma.

### GCs act directly on tumor cells to stimulate anti-melanoma immunity

Next, we sought to define the direct cellular target GCs were acting on to elicit CTL-mediated tumor control. First, we examined the contribution of GC signaling using two mouse strains in which either the cytotoxic or myeloid cell compartment lack GR expression (*Gzmb^Cre^ Nr3c1^fl/fl^* or *Lyz2^Cre^ Nr3c1^fl/fl^*, respectively) and are, therefore, selectively insensitive to GCs. GC treatment led to a significant increase in the percentage of circulating CD8^+^ T cells in wild-type and *Lyz2^Cre^ Nr3c1^fl/fl^* mice but not in *Gzmb^Cre^ Nr3c1^fl/fl^* mice (Supplementary Fig. S7A). Likewise, GC administration increased the percentage of neutrophils in wild-type and *Gzmb*^Cre^ *Nr3c1^fl/fl^* mice but not in *Lyz2^Cre^ Nr3c1^fl/fl^* mice (Supplementary Fig. S7A), confirming the specificity of GR deficiency in different immune cell types. However, tumors responded to GC administration in both strains like wild-type mice (Fig. 4A, Supplementary Fig. S7B), arguing against cytotoxic or myeloid cells being the direct target through which GCs elicit tumor control.

**Figure 4.**
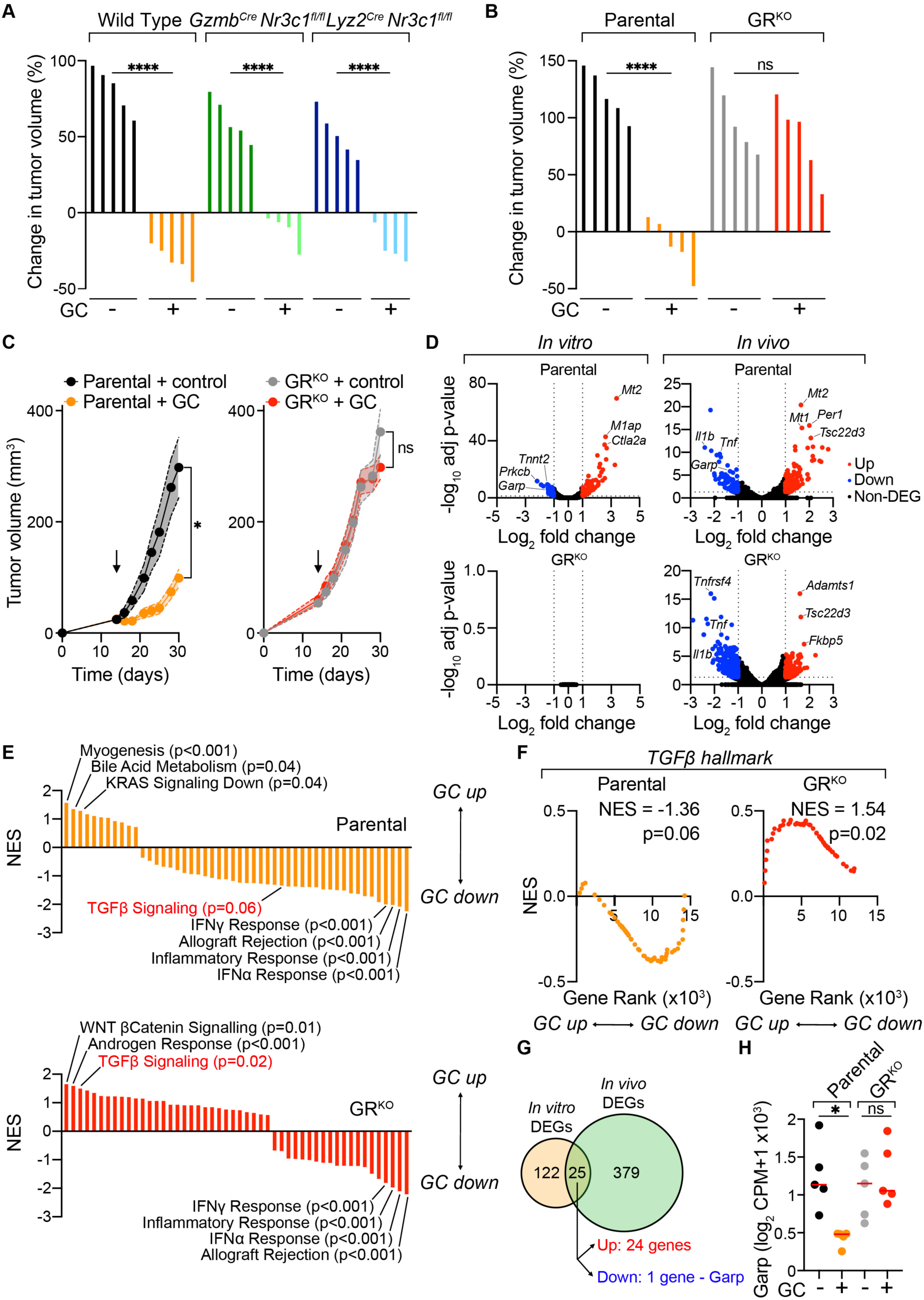
GCs act not on cytotoxic or myeloid cells, but directly on melanoma cells to trigger tumor control. (**A**) Waterfall plots showing percentage change in tumor volume at day 5 on-treatment in response to GC of melanoma tumors in wild type, *Gzmb^Cre^Nr3c1^fl/fl^* or *Lyz2^Cre^Nr3c1^fl/fl^*mice (n=4-5 per group). (**B**) Waterfall plots showing percentage change in tumor volume at day 5 on-GC-treatment of tumors formed by parental or GR^KO^ melanoma cells (n=5 per group). (**C**) *In vivo* growth profiles of parental (left) and GR^KO^ (right) melanomas treated with control or GC (n=4-5 per group). (**D-H**) RNA sequencing of parental or GR^KO^ melanoma cells *in vitro* (D, F, G), or of parental or GR^KO^ tumors *in vivo* (D-G), with or without GC-treatment (n=4-5 mice per group). (D) Volcano plots showing differentially expressed genes (DEGs) in parental (upper) or GR^KO^ (lower) cells *in vitro* (left) or *in vivo* (right). (E) Gene-set enrichment analysis (GSEA) of parental (upper panel) or GR^KO^ tumors (lower panel) showing Normalized Enrichment Scores (NES). (F) Enrichment analysis for hallmark TGF-β signaling gene set in parental (left) and GRKO (right) tumors. (G) Venn diagram comparing DEGs *in vitro* and *in vivo* in parental cells and tumors after GC treatment (from D). (H) Log2 Counts per Million (CPM) +1 Garp mRNA levels in parental or GR^KO^ melanomas. Data are expressed as mean ± SEM; one-way ANOVA (A, B, H) or two-way ANOVA (C). *, *P* < 0.05; ****, *P* < 0.0001; ns, not significant. Arrow indicates start of treatment.

We then examined whether GCs were instead signaling on melanoma cells using GR-deficient (GR^KO^) 20967 cells generated with CRISPR-Cas9 technology (Supplementary Fig. S8A, B). Crucially, tumors formed by GR^KO^ 20967 melanoma cells were fully unresponsive to GC-treatment indicating GCs act directly on tumor cells to trigger immune-dependent tumor control (Fig. 4B, C). Post-GC treatment, mice bearing GR^KO^ tumors showed a comparable reduction in spleen size and total blood CD45^+^ cells to mice bearing parental tumors (Supplementary Fig. S8C, D). These data demonstrate that GCs signal through cancer cells to drive tumor control and suggest that the broad systemic immune alterations induced by GCs cannot by themselves explain the CTL-dependent tumor suppression induced by GC treatment.

Given these findings, we hypothesized that transcriptomic analysis of control or GC-treated wild-type and GR^KO^ tumors *in vitro* and *in vivo* could uncover candidate signaling pathways and/or molecular mediators causally involved in the immune-mediated control elicited by GC treatment. RNA sequencing of cancer cells treated with GC *in vitro* revealed numerous differentially expressed genes (DEGs), including canonical GC-responsive genes in parental, but not GR^KO^, cells (Fig. 4D), demonstrating the effects of GCs on 20967 cells are mediated via GR signaling. GCs also upregulated GC-responsive genes and reduced the expression of a wide variety of inflammatory factors (such as *Il1b* and *Tnf*) *in vivo* (Fig. 4D). Unsupervised hierarchical clustering of control or GC-treated parental or GR^KO^ tumors *in vivo* identified two major clusters, one comprising all GC-treated parental tumors and the other encompassing the samples from the remaining three groups (Supplementary Fig. S8E), indicating that the effects of GCs on the bulk tumor transcriptome were largely dictated by GR signaling on cancer cells. Nonetheless, GSEA comparison of vehicle versus GC-treated tumors revealed significant downregulation of Hallmark Gene sets ‘IFNγ Response’, ‘IFNα Response’, ‘Inflammatory Response’ and ‘Allograft Rejection’ in both parental and GR^KO^ tumors post-GC treatment (Fig. 4E). These data are consistent with an overall anti-inflammatory effect of GC-treatment within tumors, regardless of whether the cancer cells are directly sensitive to GCs.

### GCs inhibit TGF-β signaling in tumor infiltrating CTLs through downregulation of melanoma cell-intrinsic GARP expression

Gene set enrichment analysis revealed ‘TGF-β Signaling’ was among the few gene-sets that were reduced by GC treatment in parental tumors but upregulated in GC-treated GR^KO^ tumors (Fig. 4E, F, Supplementary Table S3 and S4). In agreement, of the DEGs between parental and GR^KO^ tumors, many genes belonged to the ‘TGF-β signaling’ hallmark, and were downregulated in parental tumors following GC treatment (Supplementary Fig. S8E). This suggested the TGF-β pathway as a candidate signaling axis through which GCs unleash CTL-dependent tumor shrinkage. This proposed model is consistent with the multifaceted immunosuppressive functions of TGF-β in cancer immunity [22, 29–31], and particularly with its direct inhibitory effects on anti-tumor CD8^+^ T cells [32].

We thus looked for potential links to the TGF-β pathway in the DEGs identified after GC treatment of cancer cells. We reasoned that, since GCs acted directly on tumor cells to elicit their immune-mediated anti-tumor effect, any potential downstream target should be modulated by GC treatment both *in vitro* (without immune cells present) and *in vivo* (with immune cells present; Fig. 4D). Comparing DEGs between these two groups revealed 25 genes in common, 24 upregulated and only one downregulated (Fig. 4G, Supplementary Table S1). The latter was Leucine Rich Repeat Containing Protein 32 (*Lrrc32*), which encodes for Glycoprotein A Repetitions Predominant (GARP). GARP is a cell-membrane protein involved in the activation of TGF-β from its inactive, Latency-Associated Peptide (LAP)-bound form [33] and is critical for paracrine TGF-β signaling [34]. GARP expression has been reported in platelets, endothelial cells, fibroblasts and predominantly in Tregs [35–37], where it contributes to their immunosuppressive function [36, 38]. Of note, upregulation of GARP has also been reported in some cancers [39–42], including melanoma, and implicated in T cell inhibition *in vitro* [39]. We therefore hypothesized that GARP downregulation following GC treatment could diminish TGF-β activation on the surface of cancer cells, in turn releasing tumor-infiltrating CTLs from its inhibitory activity. Consistent with this hypothesis, GCs inhibited *Lrrc32* mRNA levels in parental but not GR^KO^ tumors or cells (Fig. 4D, H, Supplementary Fig. S9A), and did so within 4 hours suggesting GARP-downregulation is a direct transcriptional effect of GCs (Supplementary Fig. S9B). Moreover, analysis of previously published ChIP-seq studies revealed multiple GR binding sites in the promoter of both the murine and human GARP genes (Supplementary Fig. S9C, D).

### GARP downregulation on cancer cells, and inhibition of TGF-β signaling, are essential for the tumor-restraining effects of GCs

We then sought to assess whether GARP was mechanistically involved in the anti-tumor immune response elicited by GC treatment. For this, we first generated GARP-deficient (GARP^KO^) 20967 melanoma cells which showed no proliferative defects *in vitro* and formed comparably progressive tumors as the control GARP-expressing cancer cells *in vivo* (Fig. 5A-C, Supplementary Fig. S9E, F). Crucially, melanoma cell-intrinsic GARP-deficiency rendered tumors unresponsive to GC treatment (Fig. 5B, C). These results demonstrate that among the pleiotropic effects of GCs on the inflammatory landscape of tumors, GARP downregulation by cancer cells is indispensable for the immune-dependent tumor-inhibitory effects of GCs.

**Figure 5.**
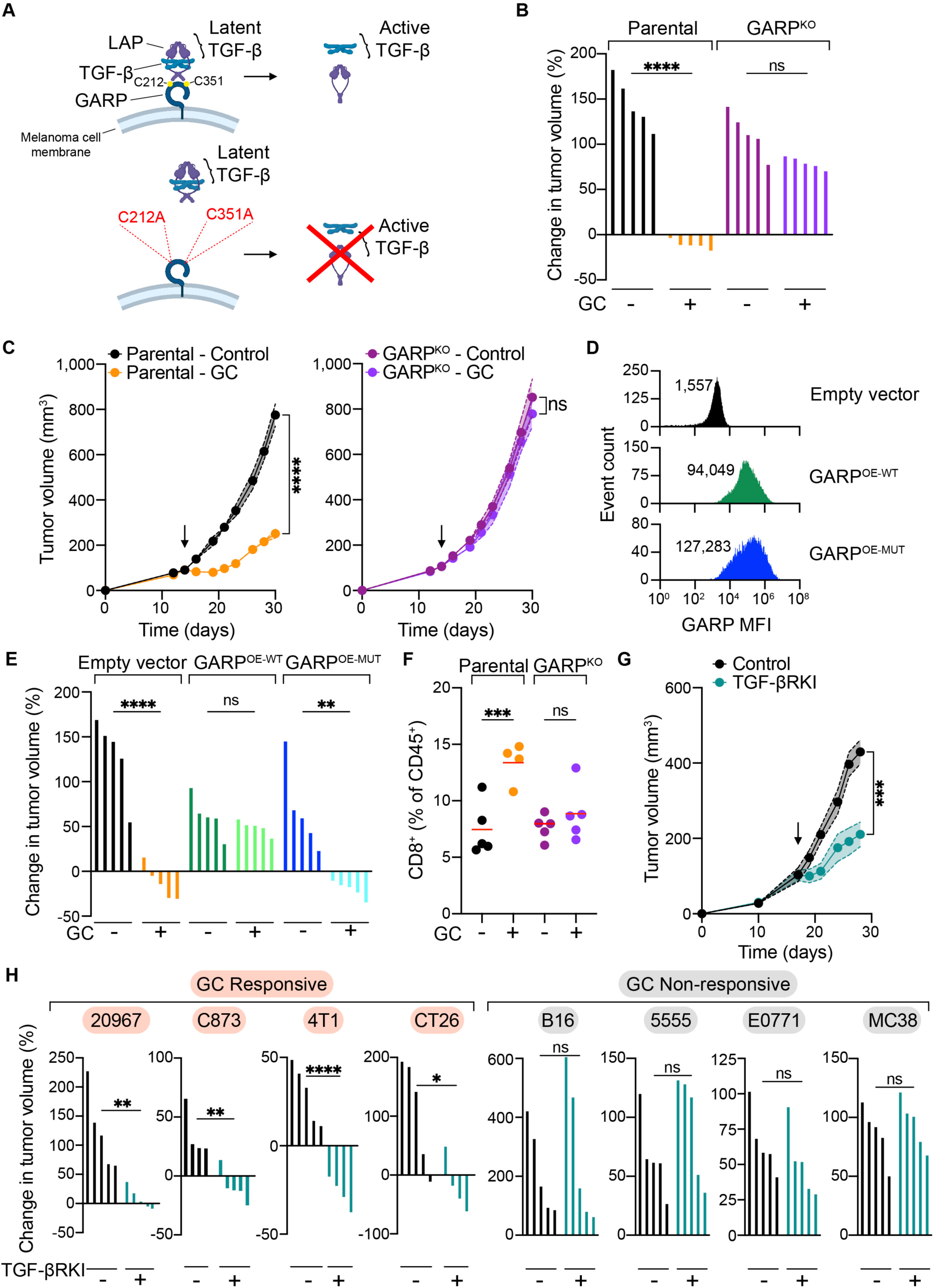
GCs inhibit the GARP/TGF-β axis in cancer cells to induce tumor control. (**A**) Function of GARP in TGFβ activation, and sites of targeted mutagenesis. (**B**) Waterfall plots showing percentage change in tumor volume of parental or GARP^KO^ melanomas at day 5 on-treatment in response to GC (n=5 per group). (**C**) Growth profiles of parental (left panel) or GARP^KO^ (right panel) melanomas treated with control or GCs (n=5 per group). (**D**) FACS analysis of surface GARP expression on empty vector, GARP^OE-WT^ and GARP^OE-MUT^ cell lines. The Mean Fluorescent Intensity (MFI) is shown. (**E**) Waterfall plots showing percentage change in tumor volume on day 5 of treatment of control and GC-treated empty vector, GARP^OE-WT^ and GARP^OE-MUT^ melanomas (n=5 per group). (**F**) Intratumoral immune-infiltrate analysis of 20967 and 20967 GARP^KO^ melanomas on day 5 post-treatment with control or GC (n=4-5 per group). (**G**) Growth profiles of 20967 tumors treated with a TGF-β receptor kinase inhibitor (TGF-βRKI) (n=5 per group). (**H**) Waterfall plot showing percentage change in tumor volume at day 5 on-treatment with control and TGF-βRKI of GC-responsive (left) or GC-non-responsive (right) tumors tumors (n=4-5 per group). Data are expressed as mean ± SEM; one-way ANOVA (B, E, F), two-way ANOVA (C, G), or unpaired t-test (H). *, P < 0.05; **, *P* < 0.01; ***, *P* < 0.001; ****, *P* < 0.0001; ns, not significant. Arrow indicates start of treatment.

We reasoned that if GARP downregulation was vital to drive tumor control following GC treatment, ectopic expression of a GC-insensitive form of GARP should render tumors resistant to GC administration. In keeping with this hypothesis, overexpression of GARP in 20967 melanoma cells (Fig. 5D) using a retroviral construct where GARP expression is driven by an unrelated constitutive promoter abrogated the inhibitory effect of GC treatment on tumor growth (Fig. 5E, Supplementary Fig. S10A). In contrast, GCs were still able to inhibit tumor growth when melanoma cells were transduced instead with a GARP mutant deficient in two cysteine residues that, based on the crystal structure of GARP [38], are critical for its binding to latent TGF-β (Fig. 5A, D, E, Supplementary Fig. S10A). These findings were confirmed in the 4T1 tumor model (Supplementary Fig. S10B) and are consistent with a previous study where over-expression of GARP increased the aggressiveness of 4T1 tumors [42]. Altogether, these results indicated the TGF-β-activating function of GARP is critical in the response to GCs. Notably, the GC-driven enrichment in tumor-infiltrating CD8^+^ T cells was not observed in GARP^KO^ tumors (Fig. 5F), consistent with the conclusion that GARP is a non-redundant critical mediator of the CTL-dependent tumor control induced by GC treatment.

We next investigated whether differential reliance on the TGF-β signaling pathway for immune evasion could account for the variable responsiveness to GCs observed across cancer models (Fig. 2E, F). To test this, we treated mice bearing GC-responsive 20967, C873, 4T1 or CT26, or GC-unresponsive 5555, B16, E0771 and MC38 models with a TGF-β receptor kinase inhibitor (TGF-βRKI). Remarkably, TGF-βRKI treatment inhibited tumor growth exclusively in cancer models that benefitted from GC administration (Fig. 5G, H). This striking concordance between tumor responsiveness to GC treatment and TGF-β signaling inhibition further supports a model in which GC-driven downregulation of GARP expression in cancer cells reduces TGF-β activation and signaling in the tumor microenvironment, thereby unleashing rapid CTL-dependent tumor growth inhibition. Furthermore, GARP^KO^ 20967 tumors no longer responded to TGF-βRKI suggesting GARP expression by 20967 cancer cells as a predominant mechanism for TGF-β activation within the tumor microenvironment (Supplementary Fig. S10C).

### GARP expression is negatively prognostic in human melanoma

To further address the human relevance of our findings, we examined GARP expression in tumor sections from an in-house melanoma patient cohort. GARP staining was heterogeneous, observed primarily within melanoma, and not immune or stromal cells (Fig. 6A, Supplementary Fig. S11A, B). Moreover, in keeping with a tumor-promoting function of GARP in melanoma cells, GARP^high^ patients exhibited a significantly worse survival in this in-house patient cohort (Fig. 6B), as well as in TCGA melanoma patients (Supplementary Fig. S12A). To specifically assess the predictive value of GARP in outcome from immunotherapy we analyzed a publicly available database of ∼1,000 ICB-treated patients [43]. Intratumoral pre-treatment GARP mRNA levels were significantly associated with worse survival (Fig. 6C). Together, these findings are in agreement with our preclinical data uncovering cancer cell-intrinsic GARP expression as an immune evasive mechanism in ICB-unresponsive tumors and pinpoint GARP as a putative biomarker of outcome and response to immunotherapy.

**Figure 6.**
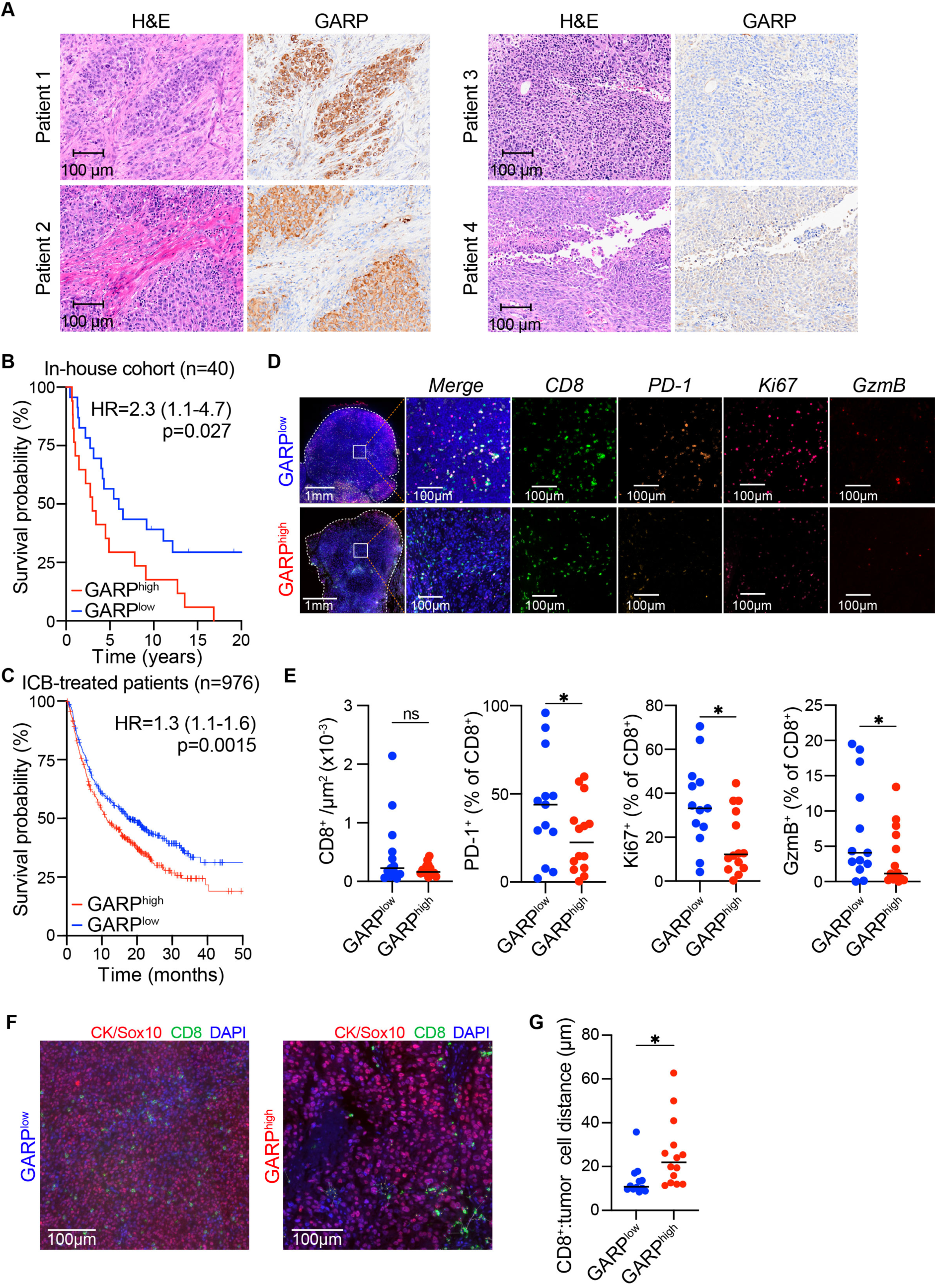
GARP expression in melanoma cells correlates with worse outcome in patients and associates with a dampened CD8^+^ T cell activation phenotype. (**A**) Representative H&E (left) and GARP (right) staining of patients with high GARP expression (Patients 1&2) and low GARP expression (Patients 3&4). (**B**) Kaplan-Meier survival analysis of 40 in-house melanoma patients stratified in GARP^high^ or GARP^low^ based on staining in (A). (**C**) Kaplan-Meier survival plot of pan-cancer *α*PD-1, *α*PD-L1 or *α*CTLA-4-treated pan-cancer patients (n=976) stratified by median GARP expression. (**D-G**) Analysis of in-house melanoma samples (B) by multiparametric immunofluorescence. Representative images (D) and prevalence of indicated populations (E) shown. Spatial analysis of distance between CD8^+^ T cells and tumor cells (F, G). Data are expressed as mean; hazard ratio (95% confidence interval), log-rank (Mantel-Cox) test (B, C) or one-tailed unpaired t-test (D, G). *, P < 0.05; ns, not significant.

We then analyzed the immune infiltrate in GARP^high^ versus GARP^low^ tumors using a 12-plex immunofluorescence panel. CD8^+^ T cells, CD4^+^ T cells, Tregs, B cells and macrophages were similarly abundant in both groups (Fig. 6D, E, Supplementary Fig. S12B, C). However, GARP^high^ tumors had significantly fewer proliferating (Ki-67^+^), Granzyme B^+^ and PD-1^+^ CD8^+^ T cells (Fig. 6D, E), consistent with an inhibitory role of GARP on CD8^+^ T cells. Moreover, spatial analysis showed CD8^+^ T cells were farther from tumor cells in GARP^high^ tumors (Fig. 6F, G). Together, these results support our preclinical data and suggest that GARP^high^ tumors are associated with poorer prognosis due to impaired CD8^+^ T cell activity.

To more directly evaluate the link between melanoma cell-intrinsic GC signaling and TGF-β activity in CTLs in humans, we next analyzed a single cell RNA sequencing dataset with available transcriptomic data for both tumor and tumor-infiltrating cells from multiple melanoma patients [44] (Fig. 7A). Among the most prevalent tumor-infiltrating immune cell populations, CD8^+^ T cells displayed the highest level of TGF-β signaling suggesting their particular sensitivity to this immunosuppressive cytokine within the tumor microenvironment (Fig. 7B, C). Given our mouse findings, we hypothesized patients with higher GR levels in melanoma cells would display lower TGF-β signaling in CTLs. Indeed, we found a pronounced and statistically significant inverse correlation between melanoma cell-intrinsic GR transcript levels and TGF-β signaling within the corresponding tumor-infiltrating CTLs (Fig. 7D). Of note, this negative association was absent or less prominent for other tumor-infiltrating immune cells (Fig. 7E). In keeping with these data, single cell RNAseq analysis of 20967 tumor-infiltrating CD8^+^ T cells showed significantly reduced TGF-β signaling activity following GC treatment (Fig. 7F). Together, these results provide further evidence to our conclusion that GCs, acting on GR expressed by melanoma cells, inhibit the GARP/TGF-β axis to enhance CTL-mediated tumor cell killing.

**Figure 7.**
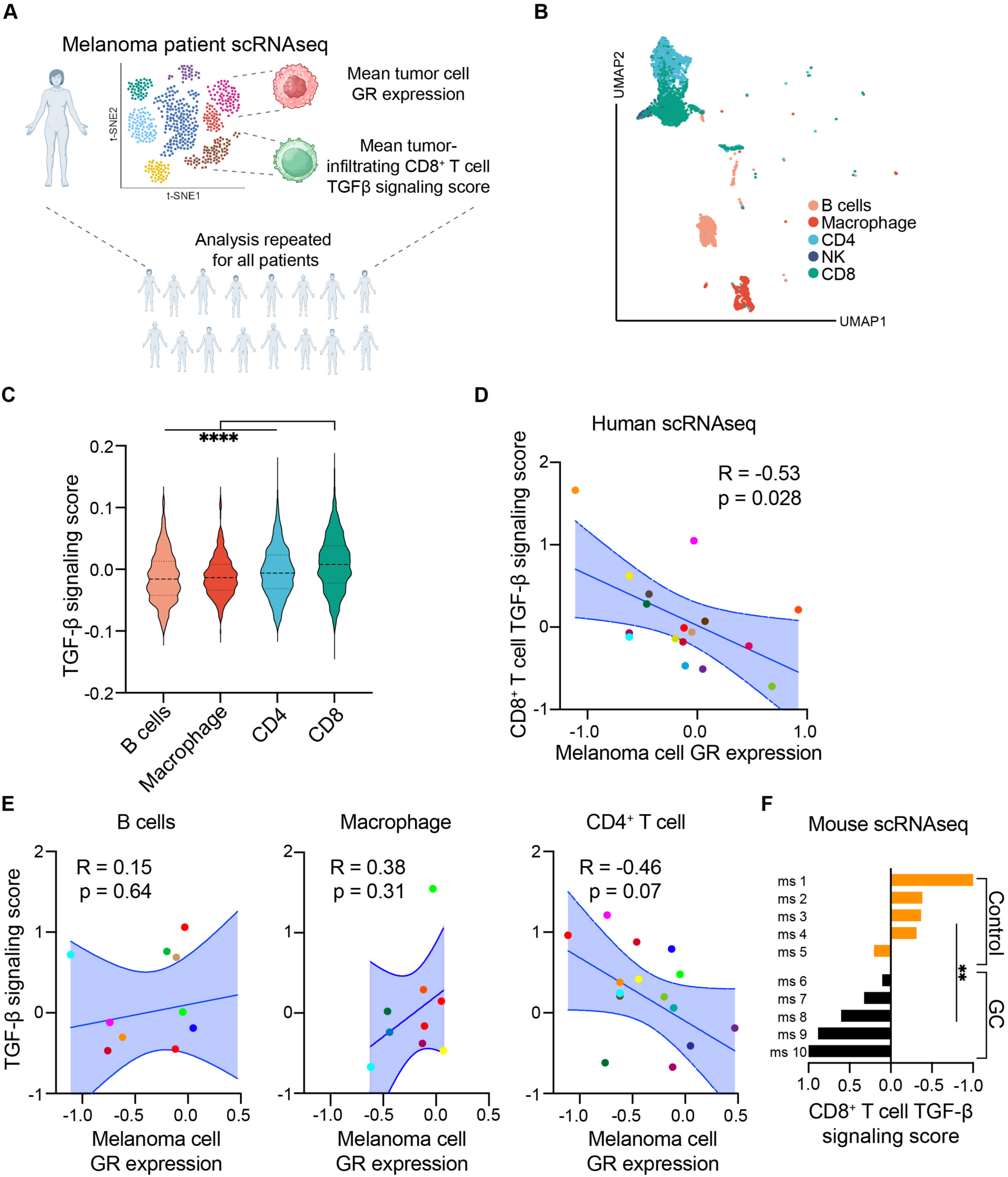
GR expression in melanoma cells anticorrelates with TGF-β signaling in tumor-infiltrating CD8^+^ T cells. (**A**) Schema showing method for determining correlation of GR expression on melanoma cells with TGF-β signaling score on tumor-infiltrating CD8^+^ T cells in patients (n=17) from Jerby-Arnon et al [44]. (**B, C**) UMAP (B) and violin plot of hallmark TGF-β signaling score (C) of different immune cell clusters from Jerby-Arnon et al [44]. (**D, E**) Correlation of GR expression on melanoma cells with TGF-β signaling score on tumor-infiltrating CD8^+^ T cells in patients (n=17; each color represents a different patient; D), and B cells, macrophages and CD4^+^ T cells (E) from Jerby-Arnon et al [44]. (**F**) Mean hallmark “TGF-β Signaling” in tumor-infiltrating CD8^+^ T cells from scRNA-sequencing analysis of control and GC-treated 20967 tumors (see Figure 3C). ms; mouse. Data are expressed as mean ± interquartile range; one-way ANOVA (C), linear regression with Pearson correlation (D-E) or unpaired t-test (F). **, *P* < 0.01; ****, *P* < 0.0001; ns.

## Discussion

Improved understanding of why large proportions of patients do not respond to immunotherapy is urgently required to improve treatment outcomes and survival. Here we show that several, but specific, murine cancer models make use of a GC-sensitive, cancer cell-intrinsic immune evasive pathway to hinder CTL-dependent anti-tumor immunity. In agreement with these findings, we show that in melanoma patients, high GR expression and GC signaling at the tumor site associate with improved survival, particularly in tumors with elevated CD8^+^ T cell-infiltration. Moreover, analysis of single cell RNA sequencing data revealed that patients with elevated melanoma cell-intrinsic GR expression have reduced levels of inhibitory TGF-β signaling in infiltrating CD8^+^ T cells. Lastly, we found that GARP levels negatively correlate with survival, including in ICB-treated patients, suggesting GARP expression may be a future therapeutic target on tumor cells or a biomarker of therapy response.

Recent work has demonstrated a benefit to combining anti-inflammatory medication, including GCs, with ICB in pre-clinical models [16–20]. A significant number of patients experience side-effects of ICB, and are often treated with immunosuppressive GCs [45]. Considerable uncertainty exists surrounding the effects of GCs on melanoma patient responses to ICB. Retrospective clinical analyses show GCs can negatively impact survival in melanoma when used early [46] or for supportive care, though GC use had no impact on overall survival when used for control of adverse events [47]. Conversely, other studies have shown that GC use in patients receiving immunotherapy has no impact on survival [48], or may even lead to improved responses [49, 50]. Our data uncovering GC signaling in melanoma patients is associated with a favorable prognosis, and that GC treatment can elicit rapid immune-dependent anti-tumor responses in select preclinical cancer models, may help explain the conflicting results of retrospective analyses of GC use in patients receiving ICB and suggest GCs may be beneficial under specific conditions.

Clarifying the role of GCs in the response to immunotherapy, and in the tumor microenvironment, is a major unanswered question in the immuno-oncology field [51]. Recently, it was shown that GCs increase pancreatic tumor cell PD-L1 expression leading to increased tumor growth [52], and that tumor cells can trigger GC production in T cells, leading to worse outcomes [53]. Additionally, poor tumor control was seen when tumor-associated myeloid cells produced GCs [54], and when tumor cells themselves produced GCs enhancing Treg activity [55]. Crucially, however, these latter three studies were conducted using B16, MC38 or E0771 cancer models, which we show here do not benefit from GC treatment, reconciling our findings with theirs. On the other hand, GCs have been shown to have a range of tumor-suppressive effects in different cancer types [56–58]. The role of GCs in the tumor microenvironment has remained controversial, and while our study does not negate that endogenous or administered GCs can be immunosuppressive and thus often tumor promoting, it provides a context in which GC signaling can be beneficial in a counterintuitive, immune-dependent, manner. Specifically, our data indicate that GCs promote immune-dependent control exclusively in tumors that are sensitive to TGF-β signaling inhibition. Accordingly, we found an absolute concordance in that, across eight independent cancer models, only those that showed impaired tumor growth post-GC treatment also did so upon pharmacological inhibition of TGF-β signaling.

TGF-β is a well-established immune-suppressive cytokine that tumors often exploit to evade immunity [29, 30, 32, 35, 59]. However, TGF-β also has the potential for anti-tumor effects, including by providing anti-growth signals, and this, together with its various pleiotropic functions in normal physiology [35, 60], has meant that the translation of anti-TGF-β therapies into the clinic is proving to be highly challenging [61, 62]. Therefore, a fuller understanding of the mechanisms underlying the biogenesis and activation of TGF-β, and of how it suppresses anti-tumor immunity, has the potential to improve anti-cancer therapies targeting TGF-β. A recent study highlighted the critical importance of the TGF-β activating complex, including latency associated peptide, GARP, and integrins, in facilitating paracrine TGF-β signaling from the GARP/TGF-β expressing cell to the receiving cell [34]. This work provides context and supports the biological relevance of our model whereby disruption of the GARP/TGF-β axis in tumor cells disinhibits directly interacting CTLs, enabling tumor cell killing. However, a potential contribution by myeloid cells or other tumor infiltrating cells to this axis cannot be excluded and warrants further investigation.

In this study, we reveal that the GARP/TGF-β axis is a cancer cell-intrinsic immune evasive pathway conserved in mice and humans that restrains the anti-tumor activity of tumor-infiltrating CTLs. We showed that in patients, elevated GR in melanoma cells is associated with reduced TGF-β signaling specifically on tumor-infiltrating CD8^+^ T cells. Moreover, high intratumoral GARP expression, detected mostly on melanoma cells, was associated with worth overall patient survival, including in patients treated with immunotherapy. This suggests context-specific targeting of the TGF-β axis, such as in GARP^high^ tumors, is likely to provide the most therapeutic benefit to patients. GARP has already been implicated in blunting cancer immunity primarily through its higher expression in Tregs [35, 36, 42, 63] and platelets [64], and its blockade improves anti-PD-1 immunotherapy responses in mice [65]. Therefore, additional uses for in-development anti-GARP therapies beyond GARP inhibition on Tregs may maximize the potential therapeutic benefit that these new drugs may have. Specifically, the understanding that this study provides – that is, that GARP can be targeted on tumour cells and not just Tregs – suggests that alternative and improved methods of patient stratification, such as those based on GARP expression levels by melanoma cells, have the potential to be an effective means of selecting those patients most likely to respond to immunotherapy, including to anti-GARP therapies.

By targeting the GARP/TGF-β axis, we show GCs can paradoxically trigger immune control in a range of select murine cancer models, even when using GCs at high doses that led to systemic anti-inflammatory and immune-suppressive effects. This provocative proof-of-concept study demonstrates that GCs can beneficially remodel the tumor immune microenvironment in specific circumstances. Human data suggests that GC signaling can be beneficial in melanoma, and that GARP represents a negative prognostic marker of overall survival and outcome from immunotherapy. The GARP/TGF-β pathway therefore represents an immunotherapeutic target on cancer cells, and new drugs selectively or locally inhibiting it could provide the beneficial anti-tumor effects of GCs without their immunosuppressive and systemic side-effects.

## Materials and Methods

### Mice

Wild-type C57BL/6 or BALB/c mice (Envigo), *Rag1^-/-^, Gzmb*^Cre^ *Nr3c1*^flox/flox^ and *Lyz2*^Cre^ *Nr3c1*^flox/flox^ mice were housed in specific pathogen-free conditions in individually watered and ventilated cages at the CRUK Manchester Institute. All procedures involving mice were performed under license from the UK Home Office granted under the Animals (Scientific Procedures) Act 1986. The CRUK Manchester Institute Animal Welfare and Ethical Review Body (AWERB) approved the overall programme of work during the Project Licence application. Tumor volumes or experimental severity did not exceed the guidelines set by the Committee of the National Cancer Research Institute [66].

### Subcutaneous 20967, 20967 GR^KO^, 20967 GARP^OE^, 20967 GARP^mut^, 5555, C873, B16, 4T1 and E0771 tumor models

For inoculation into mice, 20967, 20967 GR^KO^, 20967 GARP^OE-WT^, 20967 GARP^OE-MUT^, 5555, C873, B16, 4T1 and E0771 cells were harvested in the exponential phase of growth by trypsinization (Sigma), washed three times with ice-cold PBS (Thermo Fisher Scientific), and filtered through a 70µM cell strainer (Thermo Fisher Scientific). 1x10^5^ live cells were injected intradermally (i.d.) in 50µl of PBS into the right flank (20967, 20967 GR^KO^, 20967 GARP^OE^, 20967 GARP^mut^) or into the 4^th^ right inguinal mammary fat pad (m.f.p.) (4T1), or 2x10^5^ live cells were injected subcutaneously (s.c.) in 100µl of PBS into the right flank (5555, C873, B16, E0771) of recipient mice. Tumor growth was monitored using a hand caliper and volume was calculated by measuring the longest diameter (length) and its perpendicular (width) using the formula: (length x width^2^)/2. When tumors measured 50-200mm^3^, mice were allocated to treatment arms following normalization by tumor size across groups.

### Intravenous 20967 tumor model

For experiments assessing lung metastasis, 1x10^5^ 20967 tumor cells were injected into the tail vein following preparation as above. Mice were treated on days 5 and 7 with intraperitoneal triamcinolone. Lungs were collected on day 10, fixed in Bouin’s solution (Merck), and the number of metastases counted across the entire lung.

### Mouse Procedures

For immune cell depletion experiments, mice were injected one day before tumor cell implantation with 200μg of specific antibody intraperitoneally (IP) (anti-CD4 clone GK1.5, anti-CD8a clone YTS169.4 and anti-CD25 clone PC61, all BioXCell) and then twice-weekly with 150μg for the duration of the experiment. Depletion was confirmed via FACS. Anti-PD-1 (200µg IP injection in PBS, clone RMP1-14, BioXCell) and anti-CTLA4 (100µg IP injection in PBS, clone 9D9, BioXCell) were given twice weekly alone or in combination for 6 doses. TGF-β receptor kinase inhibitor (SB431542, Merck, 10mg/kg in 100µl of 60:40 DMSO (1 part, Sigma)/PEG400 (5 parts, Sigma):dH20) was given IP five-times weekly for the duration of the experiment. Topical cetraben cream (as control), imiquimod 5% cream, 5-fluorouracil 5% cream, diclofenac 3% cream, tacrolimus 0.1% ointment, hydrocortisone 1% cream, betamethasone valerate 0.1% cream and clobetasol propionate 0.05% ointment (all supplied by the Christie Hospital NHS Foundation Trust Pharmacy) were applied using a cotton bud to the tumor and approximately 0.5cm surrounding area three-times weekly for the duration of the experiment. Triamcinolone (25µl of 40mg/ml, or 80µl of 10mg/ml, supplied by Christie Hospital NHS Foundation Trust Pharmacy) was injected intratumorally or intraperitoneally three-times weekly for the duration of the experiment.

### FACS analysis of peripheral blood and tumor-infiltrating immune cells

For analysis of peripheral blood leukocytes, 50µl peripheral blood was taken from mice via tail-vein into an EDTA-coated capillary and transferred to 1.5ml eppendorf tubes on ice. Samples were centrifuged at 300 rcf for 6 min at 4°C to isolate plasma, FACS buffer (PBS containing 2% FCS and 0.01% sodium azide) was added to the cell pellet, which was moved to a 96-well V-bottom plate for antibody staining. Red blood cells were lysed using ACK buffer (Gibco). For analysis of tumor-infiltrating leukocytes, tumors were collected into complete RPMI on ice, cut into small pieces and digested with Collagenase IV (200 U/ml, Worthington Biochemical) and DNase I (0.2 mg/ml, Roche) for 30 min at 37°C, vortexing once. The tumors were filtered through a 70µm cell strainer and pelleted. Cell pellets were resuspended in FACS buffer. Fc receptors were blocked with anti-CD16/CD32 (clone 93, eBioscience) 5 min before staining. Cell viability was determined by Aqua LIVE/Dead-405nm staining (Invitrogen). For intracellular cytokine detection, cells were stained using the Foxp3/transcription factor staining buffer set (eBioscience) following manufacturer’s instructions. Non-specific binding of intracellular epitopes was blocked by pre-incubation of cells with 2% Normal Rat Serum (ThermoFisher). Tumor or blood samples were stained with combinations of the following antibodies: CD45-BV605 (Clone 30-F11), CD11b-BV785 or -BV421 (Clone M1/70), Ly6G-PE-CF594 (Clone 1A8), Ly6C-BV421 (Clone HK1.4), Ly6C-FITC (Clone AL-21), CD64-PE-Cy7 (Clone X54-5/7.1), F4/80-PE-Cy7 (Clone BM8), CD11c-perCP-Cy5.5 (Clone N418), CD103-PE (Clone 2E7), XCR1-APC (Clone ZET), MHCII I-A/I-E APC-eFluor780 (Clone M5/114.15.2), CD274(PD-L1)-BV421 (Clone 10F.9G2), NK1.1-PE-Cy7 or -APC (Clone PK136), CD274(PD-L1)-PE (Clone MIH5), CD3ε-PerCP-Cy5.5 or -PE-CF594 or -APC/A647 (Clone 145-2C11), CD8α-PE or -PE- Cy7 or -APC-eFluor780 (Clone 53-6.7), CD4-FITC or -APC-eFluor780 (Clone RM4- 5), CD44-APC-eFluor780 (Clone IM7), CD62L-PerCP-Cy5.5 (Clone MEL-14), PD-1-PE (Clone J43), PD-1-BV785 (Clone 29F.1A12), TIM3-APC (Clone 8B.2C12), FOXP3-PE or -FITC (Clone FKJ-16s), TCF-1-PE (Clone C63D9), TOX-APC (Clone REA473), IFNγ-eFluor450 (XMG1.2), TNF-PE-Cy7 (Clone MP6-XT22) and GARP-PE (Clone F011-5) from eBioscience, BioLegend or BD Biosciences. Following *ex vivo* restimulation for 4 hours with PMA and ionomycin, intracellular cytokines were stained using the Intracellular Fixation and Permeabilization Buffer Set (eBioscience), according to the manufacturer’s instructions. Monensin and Brefeldin A (both eBiolegend) were added two hours prior to staining. Live cell counts were calculated from the acquisition of a fixed number (5000) of 10μm latex beads (Beckman Coulter) mixed with a known volume of cell suspension. Spectral overlap was calculated using live cells or VersaComp antibody capture beads (Beckman Coulter). Cells were acquired on a Novocyte (ACEA).

### RNA isolation and Quantitative PCR (qPCR)

Tumors were collected in PureZOL Reagent (BioRad) and stored at -80°C. For processing, tumors were dissociated with 5mm Stainless steel beads (Qiagen) using the TissueLyser II (QIAGEN). Total RNA was extracted using the Direct-zol RNA Mini Prep Kit (Zymo Research) following the manufacturer’s recommendations and included a DNase digestion step. RNA was quantified using a NanoDrop One (ThermoFisher) or a Bioanalyzer (Agilent Technologies) for RNA-seq. For qPCR, total RNA was extracted from cells using RLT lysis buffer (QIAGEN) and purified using RNeasy RNA isolation kit (QIAGEN). RNA was quantified using a NanoDrop One (ThermoFisher) and 2mg cDNA was synthesized by reverse transcription using High Capacity cDNA reverse transcription kit (Applied Biosystems). qRT-PCR was performed using specific TaqMan probes (mouse *Lrrc32* FAM-MGB (assay ID Mm01273954_m1) or mouse *Hprt* (assay ID Mm03024075_m1); both Thermofisher), and TaqMan Fast Advanced Master Mix for qPCR (Applied Biosystems) using a QS5 fast real-time PCR system (Applied Biosystems). Relative quantification to the housekeeping gene *Hprt* was performed with the Δ2CT method.

### 3’-mRNA sequencing and analysis

For bulk RNA-seq, mRNA libraries were prepared using QuantSeq 3’ mRNA-Seq Library Prep Kit FWD with UDI (Lexogen) from 500 ng total RNA, quality checked by Fragment Analyzer, quantified by qPCR using the KAPA Library Quantification Kit for Illumina (Roche), and sequenced on the Illumina NovaSeq 6000. RNA-Seq reads were quality checked using MultiQC v1.4, which aggregated metrics from FastQC v 0.11.7 (https://www.bioinformatics.babraham.ac.uk/projects/fastqc/), trimmed using Trim Galore version 0.6.5, and aligned in single-end mode to the mouse genome assembly (GRCm38.75) using the STAR aligner version 2.6.1d with default parameters. Mapped data were converted to gene level integer read counts (expression) using featureCounts and Ensemble GTF annotation (Mus_musculus.GRCm38.75.gtf). Counts were filtered with minimum CPM of 0.5 in ≥3 samples and normalized using the EdgeR package of Bioconductor, and log2(CPM+1) values were used for generating heatmaps and downstream analysis. Enrichment of molecular pathways (MSigDB; [67]) was evaluated by Gene Set Enrichment Analysis (GSEA) and differential gene expression analysis was evaluated by DESeq2 using the respective GenePattern modules [68]. DEGs were defined based on linear fold change ±1.5 and adjusted *P* value/false discovery rate <0.05. Unless specified, all modules and packages were run with default parameters.

### Single cell mRNA sequencing and analysis

Tumors were processed for FACS analysis as described above and stained with Aqua LIVE/DEAD-405 nm (Invitrogen) and CD45-BV605 (Clone 30-F11) antibodies. Each tumor (n=5 per group) was stained with a TotalSeq^TM^ HTO barcode (BioLegend) prior to sorting. FACS buffer used in these experiments was EDTA and sodium azide free. Live CD45^+^ cells were sorted on a BD FACSAria III with a purity > 98%. Cells were counted using a hemocytometer after Trypan Blue exclusion. 17,100 (sample 1) or 3,500 (sample 2) live cells were loaded into a channel of a Chromium Next Gel Bead-in Emulsion (GEM) Chip G (10X Genomics, PN-1000120) and GEMs were generated on the Chromium X (10X Genomics). Indexed sequencing libraries were prepared using the Chromium Next GEM Single Cell 3ʹ Reagent Kit v3.1 (10X Genomics, PN-1000268) according to BioLegend Totalseq-A protocol and 10x Genomics user guide, with 11 cycles of cDNA amplification and 15 cycles of sample index PCR.

Libraries were quality checked by Fragment Analyzer and quantified by qPCR using a KAPA Library Quantification Kit for Illumina (Roche). Paired-end sequencing was carried out on an Illumina NovaSeq 6000 with read lengths of 28+10+10+90 cycles. Mouse single cell RNA-Sequencing (scRNA-Seq) reads were aligned and quantified with CellRanger V.7.1.0 [69] against the mm10-2020-A reference transcriptome, with introns included. Cite-Seq [70, 71] was used for ‘Cell Hashing’ to annotate cells with Hashtag Oligo (HTO) tags, enabling demultiplexing of data sets into separate samples. scRNA-Seq analysis was carried out in R 4.3.1 with Seurat V5.0.3 [72], and Scater V1.30.1 [73]. Seurat was used to demultiplex the HTO tags and identify cells associated with one HTO sample tag (singlets). Singlets were then taken for quality control. Scater identified cells of low quality, defined as an outlier by 3 median absolute deviation from the median by quality control metrics. Quality metrics included counts, genes expressed, and percentage of mitochondrial genes per cell, and were identified by Seurat. Quality controlled and filtered singlets were taken for downstream analysis in Seurat.

Library size normalization was carried out to account for sequencing depth using the NormalizeData function in the Seurat package, and 2000 highly variable genes were identified in each sample. Normalized gene counts were then scaled and Unified Manifold Approximation and Projection (UMAP) dimensionality reduction was used for visualization. Principal component analysis was completed with 50 principal components - all subsequent Seurat analyses used 50 principal components. Seurat’s Correlation Correction Analysis was used for integration, using default parameters. The resulting CCA reduction was used for clustering and UMAP generation. Cell clustering was performed by generating a shared nearest-neighbors graph, and input into a Louvain algorithm, with the resolution parameter set to 0.8. To visualize cells and clusters in a lower dimension, UMAPs were generated using default parameters. Cell-type specific markers were then identified using a Wilcoxon Rank-Sum test and used to annotate cells. The resulting filtered counts metadata information were loaded into Partek Flow software for further analysis.

The identified clusters were visualized using UMAP and different immune cell populations were classified based on group specific biomarkers (Supplementary Table S5). GSEA examining enriched Hallmark gene sets [69] in the same cell cluster from control or GC-treated tumors (Supplementary Table S6) was performed using GenePattern platform [68]. To run single-sample GSEA (ssGSEA), gene expression dataset files (.gct file), immune marker gene set file (.gmt file) and class parameter (.cls file) were uploaded on GenePattern environment and the ssGSEA score for each hallmark gene set was calculated using default parameters.

### Multiplexed immunofluorescence

Multiplexed Tyramide Signal Amplification (TSA) immunofluorescence staining was performed using the BOND RX automated platform (Leica Microsystems). 4μm sections of FFPE tumors were cut and mounted on charged slides. Dewaxing and heat induced epitope retrieval (HIER) of slides was automated on the Bond RX using ER2 (AR9640) for 20’ at 100°C. Using the Open Research Kit (DS9777), endogenous peroxidase was blocked using 3% hydrogen peroxide (VWR) for 10 minutes and the slides further blocked with 10 % w/v casein (Vector SP5020 in TBST). Antibody application, detection and TSA amplification was conducted in sequential rounds following the same general procedure: Incubation with the primary antibody in Bond antibody diluent (AR9352) for 30 minutes (in the following sequence: CD8 5μg/ml (eBioscience, 14-0808), CD4 5μg/ml (eBioscience, 14-9766) and Foxp3 2.5μg/ml (eBioscience, 14-5773-82), followed by detection using anti-rat ImmPRESS HRP (Vector MP5444) (RTU) for 30 minutes, followed by premixed TSA reagent (Perkin Elmer) 1/200 for 10’. Antibody sequence and TSA-fluorophore selection were optimized to reduce non-specific staining and tyramide binding site competition. Following labelling with TSA, each antibody was removed using a heat stripping step (epitope solution 1 (AR9961) for 10’ at 100°). Finally, nuclei were counterstained with DAPI (Thermo Fisher, 62248) for 15’ (0.33μg/ml) and mounted in coverslips with ProLong Gold antifade mountant (Thermo Fisher, P36930). Images were scanned at 20X on an Aperio VERSA (Leica Biosystems), then analyzed and quantified using the HALO® (Indica Labs) Highplex FL module.

Human hi-plex immunofluorescence was performed using a 12-plex Ultivue panel (Vizgen) on a Leica Bond platform. The antibodies used were CD20 (L26), Granzyme-B (EPR8260), CD56 (3H15L12), CD45RO (UCHL1), CD8 (C8/144b), PD- 1 (CAL20), PD-L1 (73-10), CD68 (KP1), Ki67 (SP6), CD4 (SP35), FOXP3 (236A/E7), Cytokeratin / SOX10 (AE1/AE3, BC34). Following slide drying (60°, 60 mins), standard dewaxing and antigen retrieval (20 mins, ER2), all barcoded antibodies were applied simultaneously followed by pre-amplification (room temp 25mins) and amplification (room temp 90mins) steps. Following nuclear counterstaining (room temp 15mins), four complementary barcodes with spectrally distinct fluorescent probes with were added. These detected CD20 (FITC), Granzyme-B (TRITC), CD56 (Cy5) and CD45RO (Cy7). Following a PBS wash and cover slipping (ThermoFisher prolong gold) the whole slides were scanned. Following coverslip removal in PBS and loading onto the Leica Bond platform, dehybridization of the fluorescently labelled probes using exchange buffer (37° 3x 10 mins) was performed followed by hybridization of four further complementary barcoded probes for the 2nd round of imaging. These detected CD8 (FITC), PD-1 (TRITC), PD-L1 (Cy5) and CD68 (Cy7). This was repeated a third time with the final set of four further complementary barcoded probes, following a second dehybridization step. The third round detected Ki67 (FITC), CD4 (TRITC), FOXP3 (Cy5) and Cytokeratin / SOX10 (Cy7). The Olympus VS120-L100-W-12 (Olympus Corporation, Tokyo, Japan) was utilised to image under fluorescence illumination using a Lumencor SOLA LED light source and a fast filter wheel with the Olympus penta AHF-SPX-QSEM (DAPI, FITC, TRITC, Cy5, & Cy7) filter set. Three cycles of UltiVUE imaging were aligned and fused into a single multichannel .TIF image using Indica Labs Halo 4.0.5107.488, the brightfield IHC image was deconvolved to isolate stained areas using the Deconvolution (1.1.10) module and subsequent output was aligned to the .TIF image and fused to produce a final 14-plex image. Analysis was carried out in Halo with manual annotation for tumor areas followed by cell detection and positivity measurement using the Highplex FL v4.3.2 module. Subsequent single cell object data was taken into the Spatial Analysis and analyzed using the Nearest Neighbor method.

### Chromogenic immunohistochemistry

Slides were stained on the BOND RX automated platform (Leica Microsystems). 4μm sections of FFPE tumors were cut and mounted on charged slides. Dewaxing and heat induced epitope retrieval of slides was automated on the Bond RX, using Epitope Retrieval Solution 1 (ER1) (Leica Microsystems, AR9961) for 20 minutes at 98 °C. Using the Refine kit (Leica Microsystems, DS9800), endogenous peroxidase was blocked for 10 minutes and the slides further blocked with 10% w/v casein (Vector, SP5020) in TBST for 20 minutes. Following anti-GARP (Proteintech 26021-1-AP) antibody incubation (1hr at RT) at a concentration of 2.66μg/ml, rabbit envision (Agilent K4003) was used as detection for 30 minutes, followed by Refine kit DAB (as per manufacturer’s instructions). Slides were finally dehydrated through graded ethanols, cleared in xylene and coverslipped with Pertex. (CellPath SEA-0100-00A). Slides were scanned using a VS200 slide scanner (Olympus). Whole slide imaging was performed using the Olympus VS200 MTL (Olympus Tokyo, Japan), in conjunction with an Olympus UPLXAPO20X (NA 0.6, 0.274 μm/pixel) objective lens. The slides were viewed using QuPath v0.5.1 [74] and scored for GARP expression whilst blinded to survival data using the hot-spot H-scoring system: a 1mm x 1mm section of prominent staining in each tumor was assessed for GARP protein levels, and scored out of 300 using the following formula: %no positivity * 0 + %low positivity * 1 + %medium positivity * 2 + %high positivity * 3.

### Hematoxylin and eosin staining

Melanoma biopsies were obtained with ethical approval (including full informed consent from patients prior to sample donation) from Manchester Cancer Research Centre Biobank (MCRC Biobank Research Tissue Bank Ethics ref: 22/NW/0237). Samples were stained with hematoxylin and eosin on the Leica Autostainer XL. Following incubation at 60°C for 10 minutes, FFPE sections were dewaxed in xylene (3 x 8 minutes) and rehydrated through graded ethanol (100% x3, 90% and 70%). Following a water rinse, sections were stained with Gills 2X hematoxylin (Epredia 6765008) for 3 minutes. Sections were then blued in Scott’s tap water (Surgipath 3802901E), rinsed in water and stained with alcoholic eosin (Epredia 6766008) for 1 minute. Sections were then dehydrated through graded ethanols, cleared in xylene and coverslipped with Pertex (CellPath SEA-0100-00A).

### Cell lines and cell culture

The 20967 and C873 cell lines kindly provided by Richard Marais were derived from the C57BL/6 Nras^+LSL-G12D^;Tyr::CreERT2^+/o^ model [75]. The 5555 cell line was derived from the C57BL/6 Braf^+LSL-V600E^;Tyr::CreERT2^+/o^;p16^INK4a-/-^ model [76].

All cancer cell lines except E0771 were cultured under standard conditions in RPMI-1640 (Sigma-Aldrich) supplemented with 10% heat-inactivated fetal bovine serum (Sigma-Aldrich) and 1% Penicillin/Streptomycin (Sigma-Aldrich). E0771 cells were cultured in high-glucose DMEM (Sigma-Aldrich) supplemented with 10% heat-inactivated fetal bovine serum (Sigma-Aldrich), 1% non-essential amino acid (Gibco), 1% sodium pyruvate (Gibco), and 1% Penicillin/Streptomycin (Sigma-Aldrich). All cell lines were routinely confirmed to be mycoplasma-free (Venor® GeM gEP Mycoplasma Detection Kit, Minerva Biolabs) and mouse hepatitis virus-free (QIAamp® Viral RNA Mini extraction kit, Qiagen) by qPCR.

20967 GR^KO^ cells were generated by ribonucleoprotein (RNP)-mediated CRISPR/Cas9-mediated editing (Integrated DNA Technologies) following manufacturer’s protocol. To produce the RNP complex, 1.5pmol Cas9 enzyme was combined with 1.5pmol crRNA:trRNA duplex (Nr3c1 crRNA: 5’- CAATTTCACACTGCCACCGT-3’) in Opti-MEM media (Gibco) and incubated at room temperature for 5 min. The RNP complex was then combined with Lipofectamine CRISPRMAX (Invitrogen) and Opti-MEM and incubated at room temperature for 20 min. 20967 cells were trypsinized, washed with PBS and 2x10^4^ cells were added to the transfection mixture in a 24-well plate. 48h post-transfection 500 single cells were re-plated in 10cm dishes and incubated for 7 days to develop into macroscopic colonies. GR expression in single clones was verified by western blotting using GR specific antibody (Cell Signaling Technology).

20967 GARP^OE-WT^ overexpressing cell lines were generated by retroviral transduction of the pFB-neo vector encoding full-length cDNA of Lrrc32 cloned from 20967 WT cells. Primers used were: Fwd 5’- AATTCGGAATGAGCCACCAGATCCTGCT-3’; Rev: 5’- CTCGAGGATCAGGCTTTGTATTGTTGGCTGAG-3’. Full-length Lrrc32 cDNA was subcloned into pFB-neo vector using Gibson assembly and these primers: Fwd: 5’- AAGCCTGATCCTCGAGCGGCCG-3’; Rev: 3’- GTGGCTCATTCCGAATTCGTCGACAATTCGATC -3’.

For generating 20967 GARP^OE-MUT^, C212A and C351A mutations were generated using NebBuilder HiFi DNA Assembly and these primer pairs: Fwd C212A_C351A 5’- CTCCCTCACCgcgATCTCAGACTTC-3’; Rev C212A_C351A 5’- CAAAGGATCGCAGcgcGTTTCTGCTG-3’; Fwd C351A_C212A 5’- CAGCAGAAACgcgCTGCGATCCTTTG-3’; Rev C351A_C212A 5’- GAAGTCTGAGATcgcGGTGAGGGAG-3’. Transduced 20967 cells were selected using 300ug/ml G418.

### Treatment of cancer cells *in vitro*

5x10^3^ (96-well plate for IncuCyte experiments), 2x10^5^ (12-well plate for RNA) or 5x10^5^ cells (6-well plate for protein) were seeded overnight. The next day, the culture medium was replaced with fresh containing DMSO or clobetasol propionate and left for 48 hours. Cells were treated with 10µM clobetasol propionate, unless stated otherwise.

### IncuCyte live-cell imaging

Parental and mutant 20967 cells were seeded at 5x10^3^/well in triplicate in a clear 96-well plate and allowed to adhere to the well bottom at room temperature for 20 minutes. Imaging of cells was performed using an IncuCyte S3 imaging system (Essen BioScience). Images were taken at 10X magnification, capturing 4 fields of view per well every 2 hours for 5 days.

### Western blotting

NP40 cell lysis buffer (Invitrogen) supplemented with 1xPMSF (Sigma) and 1x complete protease inhibitor cocktail (Roche) was added directly to PBS-washed cells in a 6-well plate on ice, cells were scraped to collect lysates. Lysates were centrifuged at 15000 rpm for 20 min at 4°C and total proteins quantified using the Pierce BCA Protein Assay kit (Thermo Fisher Scientific). 10µg of protein was diluted in 1x laemmli buffer (Bio-Rad) with 2.5% β-mercaptoethanol, denatured at 95°C for 5 min and loaded onto 10% Mini-PROTEAN TGX Gels (Bio-Rad). Proteins were transferred to nitrocellulose membranes using the Trans-Blot Turbo system (Bio-Rad) and membranes blocked with Intercept PBS blocking buffer (Li-COR) for 1h at room temperature. Membranes were incubated overnight at 4°C in Li-COR blocking buffer containing 0.2% Tween-20 with primary antibodies against specific antibodies (Glucocorticoid receptor (D6H2L, Cell Signaling Technologies #12041S) or β-tubulin (D3U1W, Cell Signaling Technologies #86298S). Membranes were washed with PBS containing 0.1% Tween-20 and incubated with secondary antibodies IRDye 680RD Goat Anti-Mouse IgG and IRDye 800CW Goat Anti-Rabbit IgG (both 1:15000, Li-COR) for 1h at room temperature. Membranes were washed with PBS containing 0.1% Tween-20 again and protein bands were visualized using the Odyssey CLx system (Li-COR) and processed using the ImageStudio software (Li-COR).

### Development of GC-stimulated gene signature

*In vitro* and *in vivo* control- and GC-treated 20967 cells or tumors, respectively, underwent bulk RNA sequencing. DEGs were compared between *in vitro* and *in vivo* tumors and identified 25 overlapping genes. 24 were upregulated (Fig 3F, Supplementary Table S1), and the human homologs from this list (Supplementary Table S2) were used to create the signature.

### Transcriptomic Analysis of Patient Datasets

Survival analysis of human patients was performed on the TCGA PAN-CANCER dataset (10,952 patients) and the TCGA SKCM melanoma dataset (458 patients), with expression TPMs downloaded from the Genomic Data Commons (GDC) (https://portal.gdc.cancer.gov), and Log2+1 FPKM expression and survival data downloaded from the Xena platform [77]. The Leeds Melanoma Cohort (LMC; 703 patients) is a cohort of FFPE primary melanoma blocks, sampled using the Illumina DASL Human HT12 v4 array, as described previously [26] and is available from the European Genome-phenome Archive (accession no. EGAS00001002922). Patients were stratified into top and bottom quartile of *NR3C1*, *11BHSD1*, *11BHSD2* and *LRRC32* expression, and survival compared. Multivariate analysis was performed using a Cox proportional hazard model. Briefly, the *NR3C1* gene was treated as a continuous variable in both TCGA and LMC, and compared to the indicated covariates. For correlation of GC gene signature expression with *NR3C1* levels, TPM data was downloaded from GDC, log2 transformed, and correlated to *NR3C1* expression levels. For correlation of the GC signature with survival, the GC signature score (Supplementary Table S2) was calculated for each patient based on a sum of the relevant genes’ Log2(FPKM+1) expression data downloaded from the Xena platform and correlated with survival. For Fig 3H, patient CD8^+^ T cell infiltration (CD8inf) levels were inferred by MCP Counter [78] and downloaded from the Tumor Immune Estimation Resource [79], and patients stratified by both median CD8inf into CD8inf^high^ and CD8inf^low^ groups, and by NR3C1 top 25% and bottom 25% quartile expression. Survival analysis of ICB-treated human patients stratified by GARP expression was performed using the freely available KMplot platform [43].

### Human single cell RNA sequencing analysis

To investigate prior datasets, scRNA-Seq counts and cell annotations from Jerby-Arnon et al.^29^ were accessed from GEO (Accession: GSE115978) and loaded into R for normalization, identification of 3000 highly variable genes, dimensionality reduction and further analysis via Seurat. To assess for associations between gene expression and known signaling pathways, the KEGG TGF-β signaling pathway gene set [80] was inputted into Seurat’s AddModuleScore function. The TGF-β signaling module score was averaged per sample in the malignant cells and correlated against gene expression in CD8^+^ T cells via Pearson’s and assessed for statistical significance. Samples with fewer than 10 malignant or CD8^+^ T cells were excluded from the comparison.

### Statistical analysis

Graphs were plotted using GraphPad Prism v10.2.2 (GraphPad Software Inc.). Flow cytometry standard (.fcs) files were analyzed using FlowJo version 10.8.1. (Tree Star Inc.). IncuCyte images were analyzed using IncuCyte software GUI v2020C Rev1 (Essen BioScience). Experiments were performed at least twice independently with similar results. Statistics were calculated with GraphPad Prism or directly in R, and values expressed as mean ±SEM. Data were analyzed with the following tests (see figure legends for details): Unpaired two-tailed Student’s t-test (or one-tailed if indicated), unpaired Mann-Whitney t-test for non-Gaussian distributed data, one-way ANOVA tests adjusted for multiple comparisons using Dunnett’s test, two-way ANOVA using Tukey’s multiple comparison test, Log-rank (Mantel-Cox) test, multivariate COX regression analysis and linear regression analysis with Pearson correlation. A *p* value < 0.05 (\**p*<0.05; \*\**p*<0.01; \*\*\**p*<0.001; \*\*\*\**p*<0.0001) was considered significant.

### Schema generation and visualization

Schema in Figures 5 and 7 were created using Biorender.com. Heatmaps were generated using the ClustVis resource [81].

### Data availability

scRNA-Seq counts and cell annotations from Jerby-Arnon et al. were accessed from GEO (Accession: GSE115978). Leeds Melanoma Cohort data is available from the European Genome-phenome Archive (accession no. EGAS00001002922). TCGA data were downloaded through the Genomic Data Commons (https://portal.gdc.cancer.gov) or XenaBrowser (https://xenabrowser.net). Immunotherapy-treated patient analysis was performed on the KMPlot cohort (https://kmplot.com/analysis/). MCP Counter data was downloaded from the Tumor Immune Estimation Resource. (http://timer.comp-genomics.org/timer/).

Sequencing data are deposited in the NCBI’s Gene Expression Omnibus database and can be accessed through GEO. The accession number for bulk tumor transcriptomes of surgically excised 20967 melanoma tumors treated with control or GCs, and *in vitro* 20967 parental or GR^KO^ control or GC-treated cells, will be GSE286519. The accession number for single-cell transcriptomes of tumor-infiltrating leukocytes in 20967 melanoma tumors treated with control or GCs will be GSE286520.

## Authors’ Contributions

CHE and SZ conceived the study. CHE, PD and SC performed experiments with assistance from AM, MAK, EB, MR, LNB, KH, ER, AP, CRB, and assistance with analysis from AB, RR, RS, SS, VF, MGR, TB, JN. SZ acquired funding. CD, JN, JNB, SS, CEMG provided supervision. CHE and SZ wrote the manuscript with input from all authors.

## Acknowledgments

We thank colleagues in CRUK Manchester Institute Core Facilities (including the Histology, Biological Resources Unit, Transgenic Breeding Facility, Molecular Biology, Flow Cytometry and Visualization, Irradiation and Analysis facilities) for their assistance. Richard Marais provided the 5555, 20967 and C873 cell lines. We thank Dr Anshuman Chaturvedi for assistance with melanoma patient IHC interpretation. This work was supported by a Wellcome Trust 4ward North Clinical PhD Fellowship (CHE), Ian Nelson Doctoral Scholarship (CHE), Cancer Research UK Institute Award (C5759/A20971 and A27412 - SZ), Cancer Research UK grants C588/A19167, C8216/A6129 and C588/A10721 (JN, JNB) and a National Institutes of Health grant CA83115 (JN, JNB). Supported by the NIHR Manchester Biomedical Research Centre.

## Supplementary Figures

**Supplementary Figure 1.**
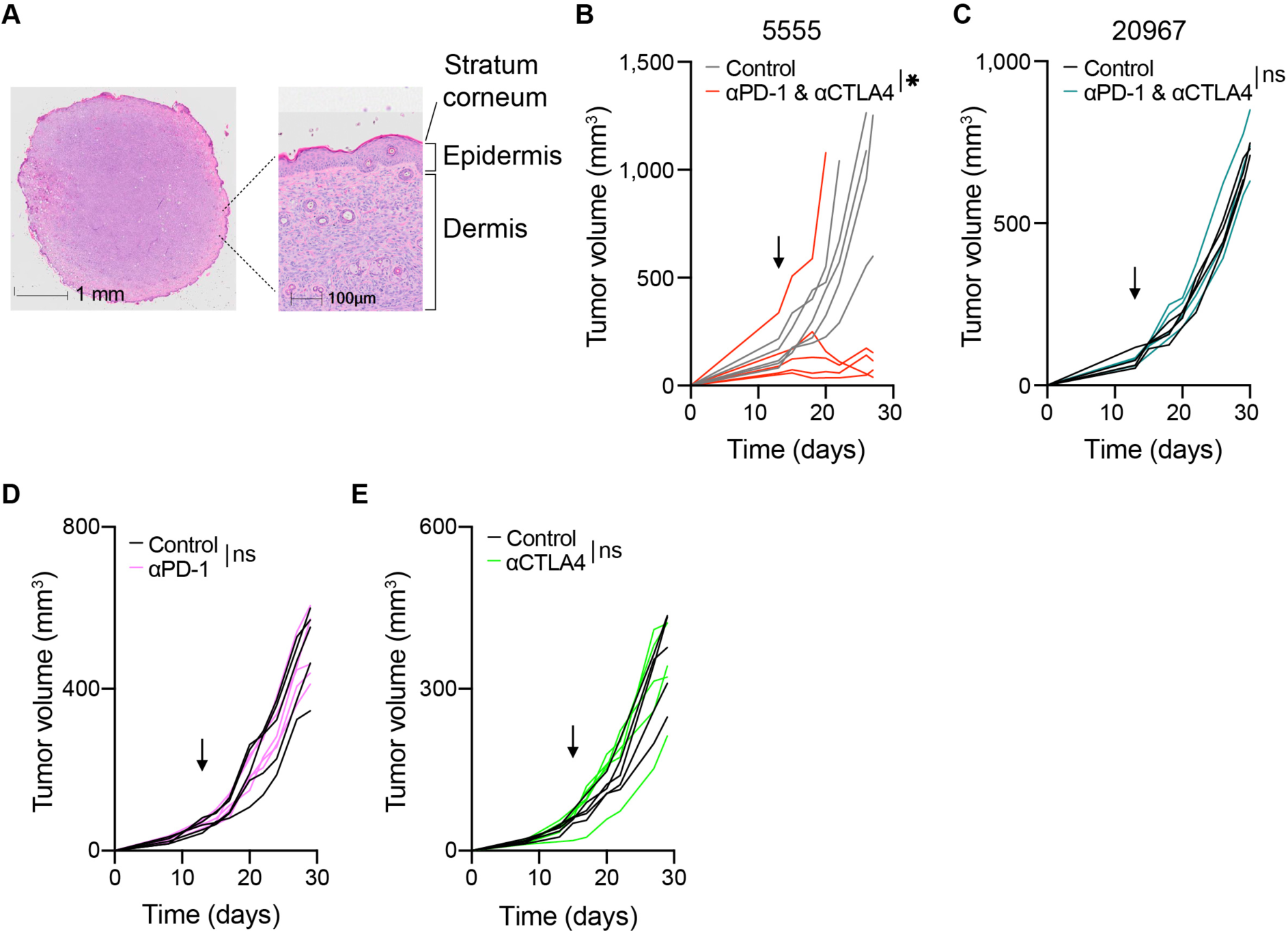
The 20967 melanoma model is resistant to immune checkpoint blockade. (**A**) Representative hematoxylin and eosin slide of an intradermal 20967 melanoma tumor. (**B-E**) Individual tumor growth profiles of 5555 (B) and 20967 tumors (C-E) after treatment with combination αPD-1 and αCTLA4 (B, C), αPD-1 only (D) or αCTLA4 only (**E**) (n=5 per group). Arrow indicates start of treatment. Two-way ANOVA (B-E). *, *P* < 0.05; ns, not significant.

**Supplementary Figure 2.**
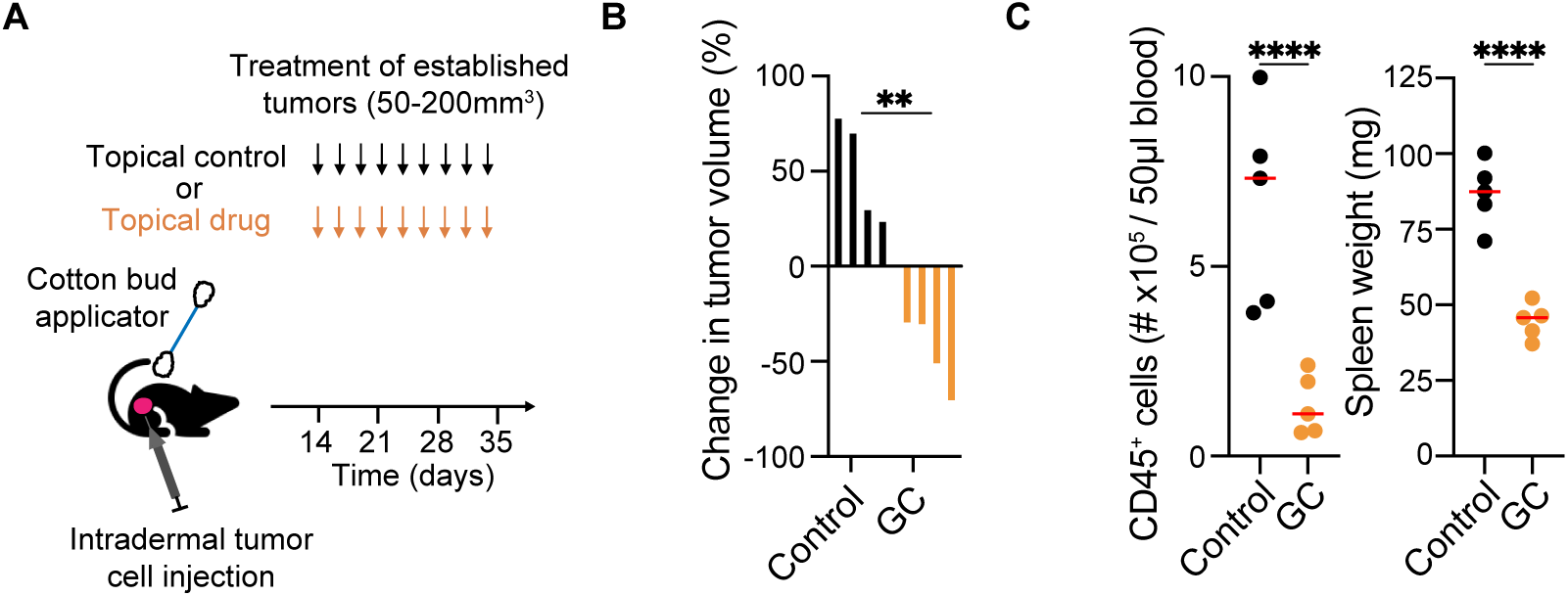
Topical GC treatment of 20967 tumors reduce leukocyte counts systemically. (**A**) Treatment schedule of melanoma with topical drug. (**B**) Waterfall plot showing percentage change in tumor volume at day 5 post-treatment of 20967 tumors over 300mm^3^ following control and GC-treatment (n=4 per group). (**C**) Peripheral blood CD45^+^ cell count on day 14 after treatment start (left panel) and spleen weight (right panel) on day 5 post treatment start with GC or control (n=5 per group). Data are expressed as mean; unpaired t-test (B, C). **, *P* < 0.01; ****, *P* < 0.0001.

**Supplementary Figure 3.**
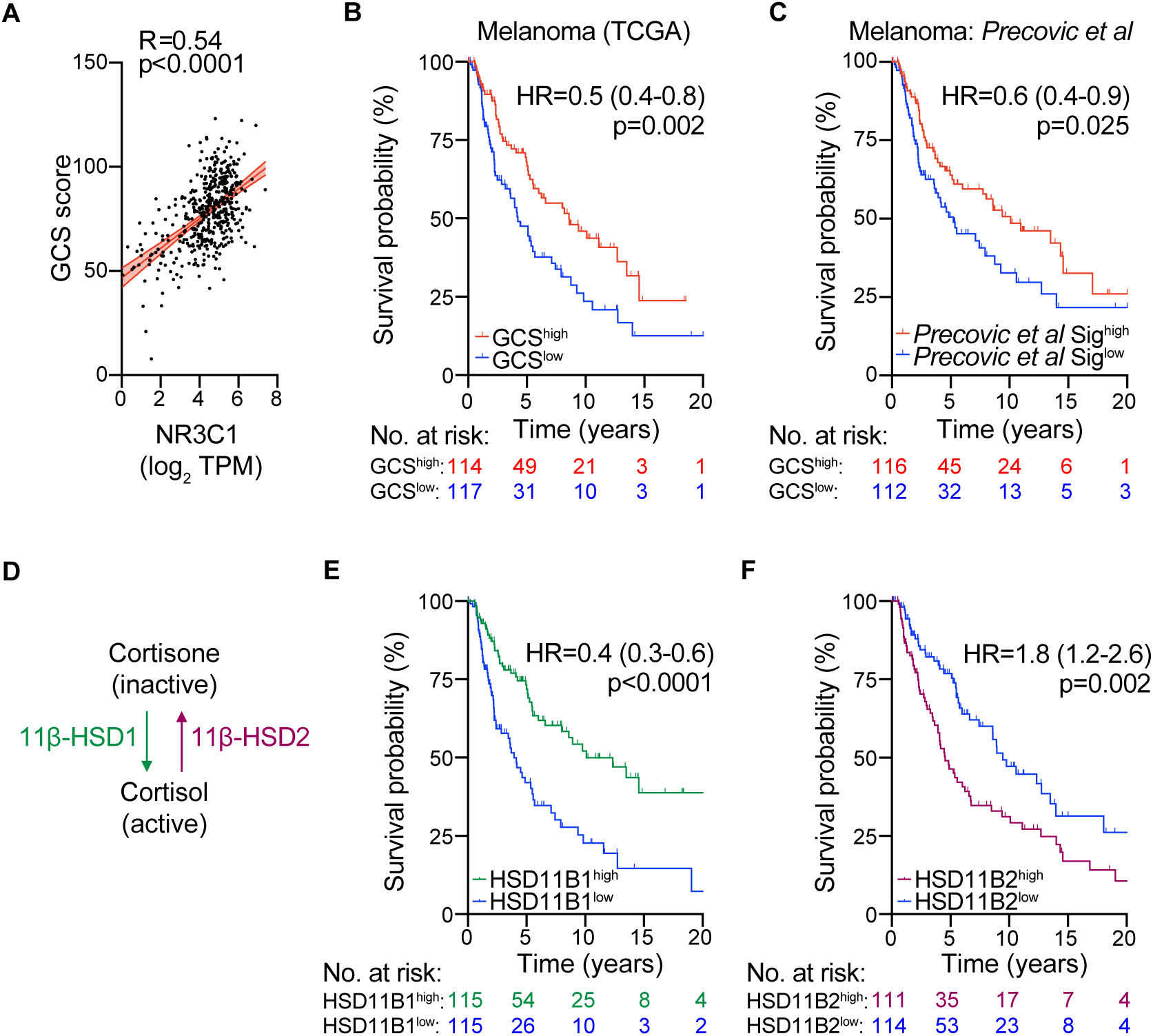
Intratumoral GC receptor expression, signaling and levels are associated with improved survival in melanoma. (**A**) Correlation, with 95% confidence interval shown, between GR expression and GC Signature in melanoma patients (TCGA, n=458). (**B, C**) KM survival analysis based on GC signature (GCS; described later, Table S2, B) or previously published GC signature (C), stratified by upper and lower quartiles, of TCGA melanoma (n=458) patients. (**D**) Pathway controlling active glucocorticoid levels in peripheral tissues. (**E, F**) KM survival plots of TCGA melanoma patients (n=458) stratified by upper and lower quartile of 11β-HSD1 (F) and 11β-HSD2 expression (G). Linear regression with Pearson correlation (A), Hazard ratio (95% confidence interval) or log-rank (Mantel-Cox) test (B, C, E, F).

**Supplementary Figure 4.**
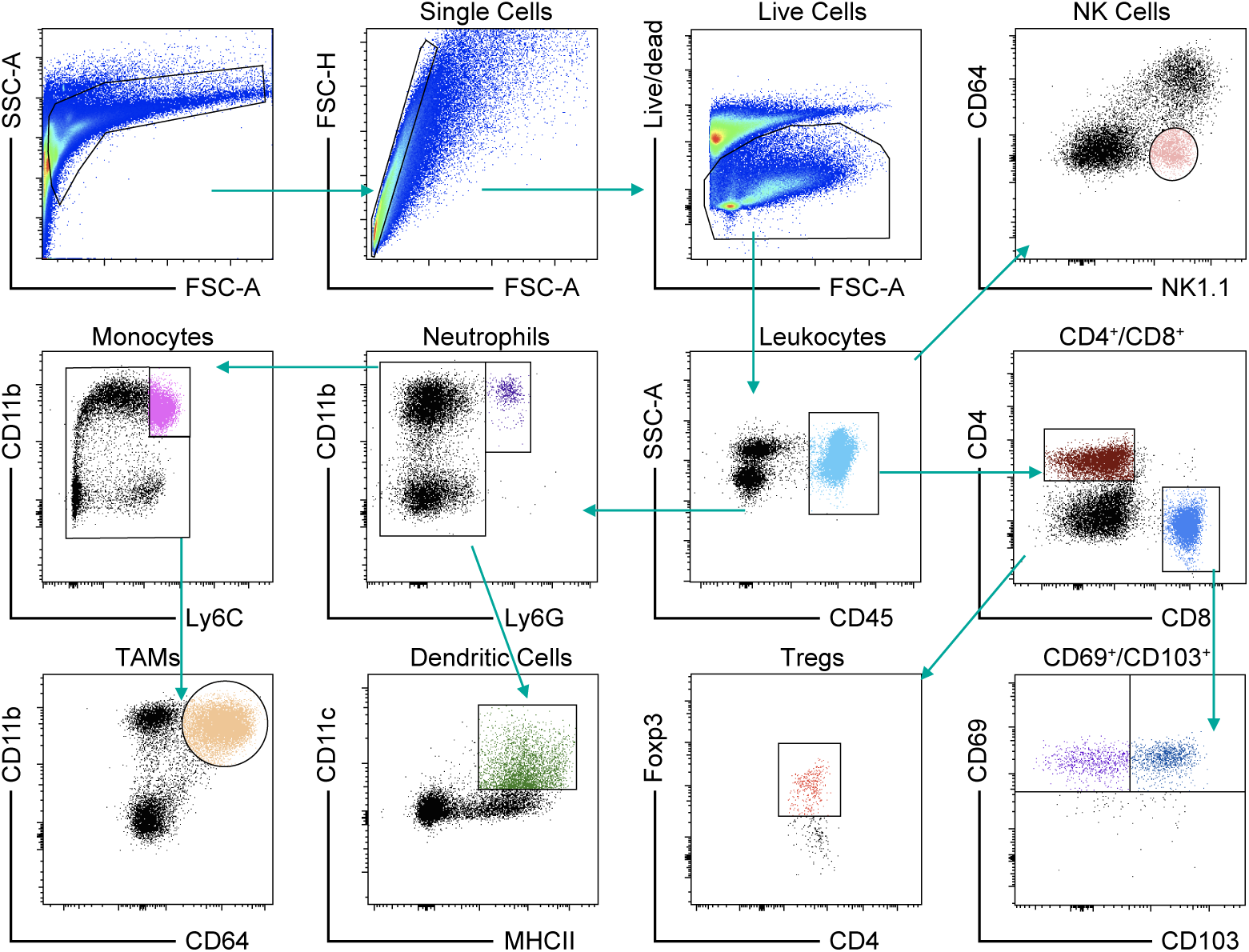
Gating strategy used for intratumoral immune infiltrate analysis. Gating strategy for intratumoral immune-infiltrate analysis of 20967 melanomas (related to Figures 2, 3 and Supplementary Figure 5).

**Supplementary Figure 5.**
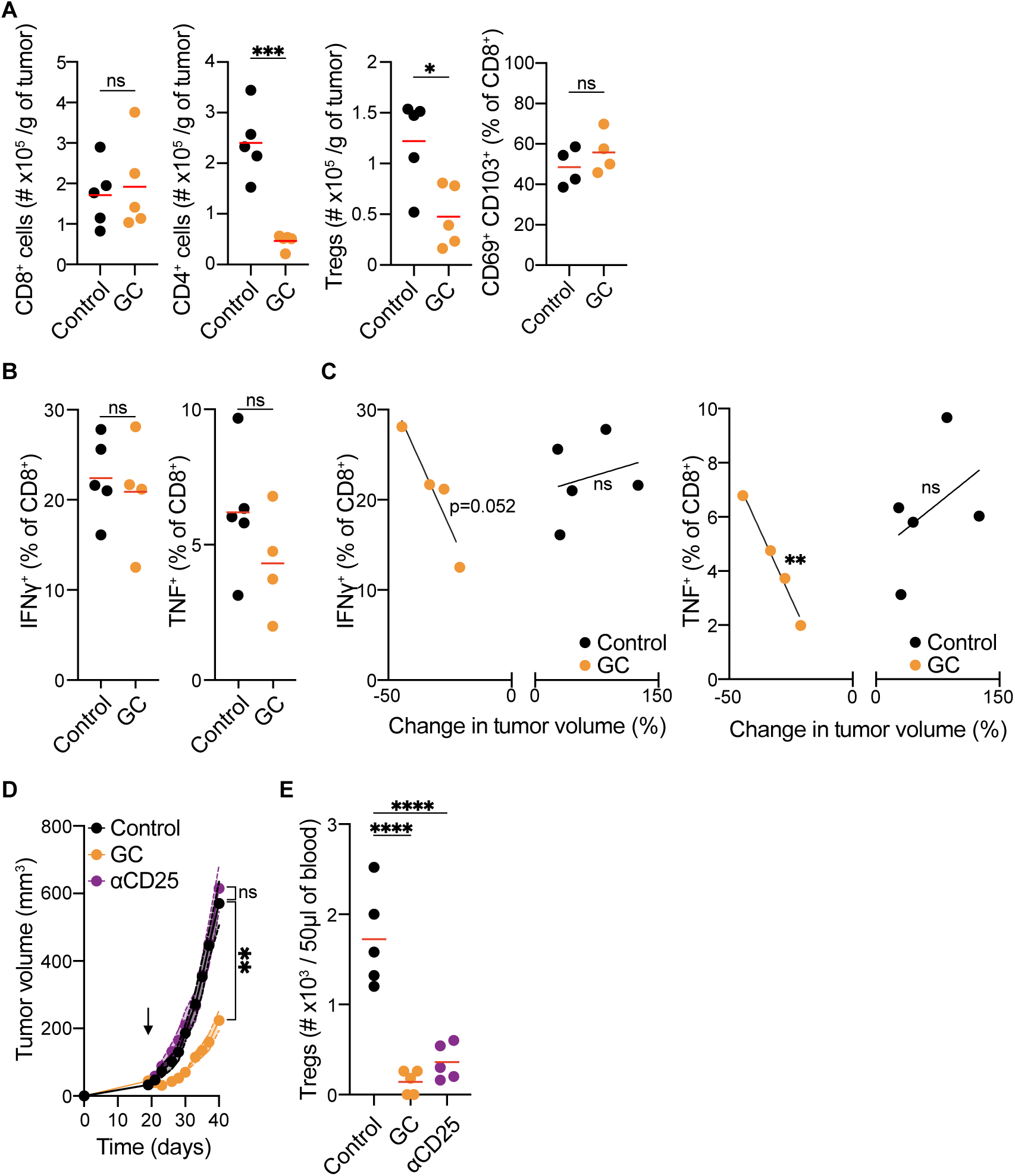
CD8^+^ T cell activity correlates with extent of GC-induced tumor shrinkage, and Treg depletion does not phenocopy GC treatment. (A) Intratumoral immune-infiltrate analysis of 20967 melanomas on day 5 post-treatment with control or GCs (n=5 per group). (**B, C**) Intracellular cytokine production analysis of CD8^+^ T cells in control or GC-treated melanomas on day 5 post-treatment (n=4-5 per group) showing proportion (B) or correlation with change in tumor volume (C). (**D, E**) 20967 tumor growth curves of GC-treated mice, or control-treated tumor-bearing mice depleted or not of Tregs (D) (n=5 per group), and peripheral blood analysis of CD4^+^ Foxp3^+^ cells (E). Arrow indicates start of treatment (D). Data are expressed as mean ± SEM; unpaired t-test (A, B), simple linear regression (C), two-way ANOVA (D) or one-way ANOVA (E). *, *P* < 0.05; **, *P* < 0.01; ***, *P* < 0.001; ****, *P* < 0.0001; ns, not significant.

**Supplementary Figure 6.**
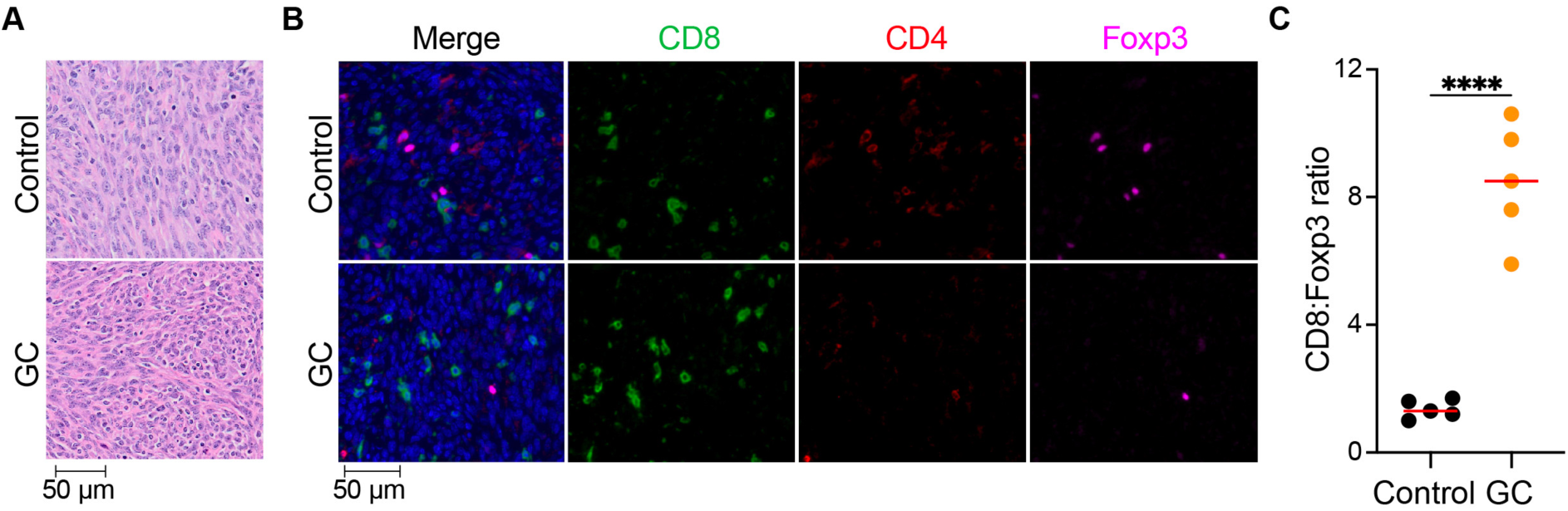
GC treatment spares intratumoral CD8^+^ T cells, resulting in an increased ratio of CD8^+^ T cells to Tregs. (**A**) High magnification Hematoxylin & Eosin images of day 5 post control or GC-treated tumors. (**B**) Representative immunofluorescence images of control and GC-treated melanomas showing individual stains used (n=5 per group). (**C**) CD8:Foxp3 ratio from immunofluorescence analysis of control and GC-treated melanomas (n=5 per group). Data are expressed as mean; unpaired t-test (C). ****, *P* < 0.0001.

**Supplementary Figure 7.**
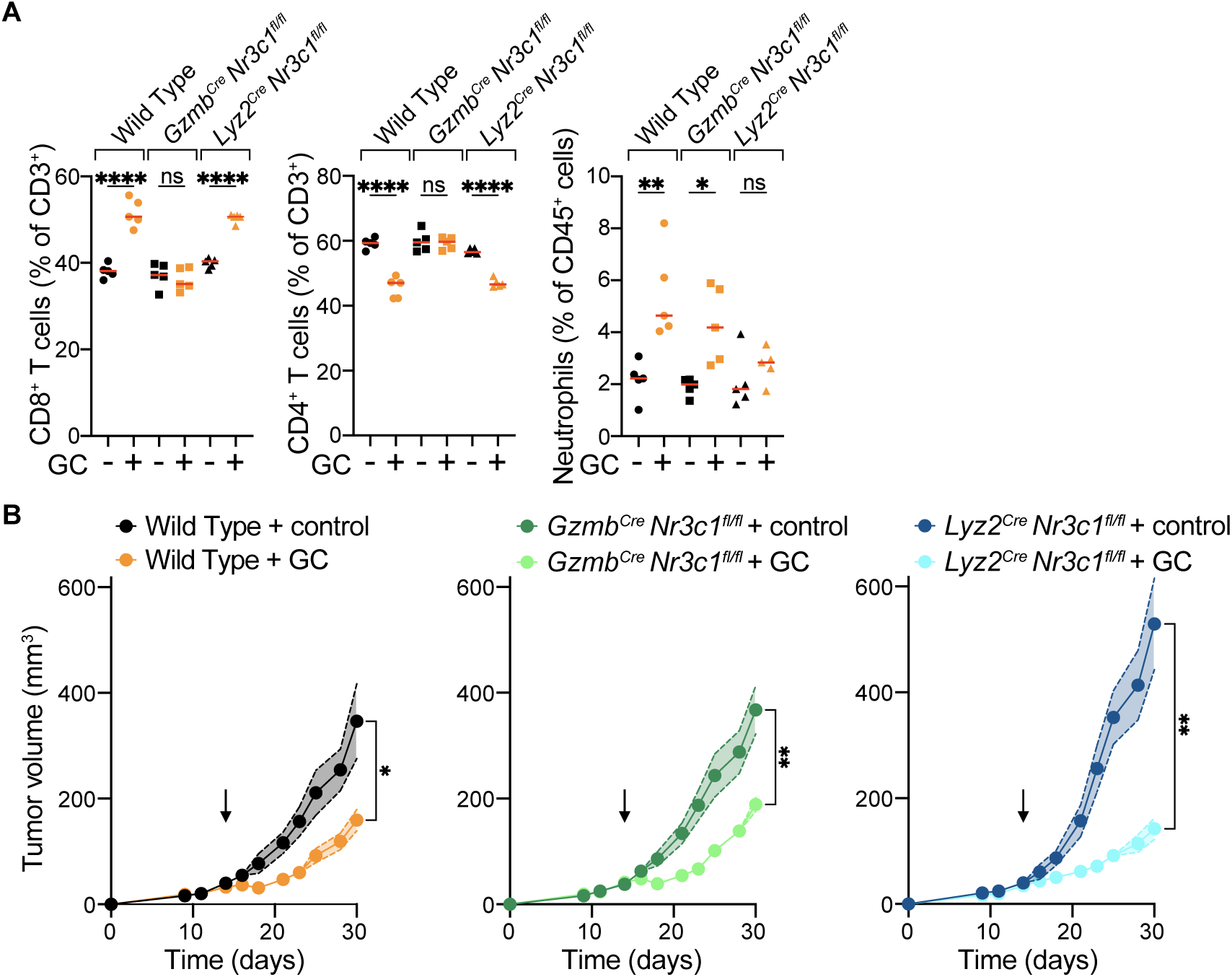
GCs do not act on immune cells to trigger tumor control. (**A**) Peripheral blood CD8^+^ T (left panel) and CD4^+^ T (middle panel) cells (as a percentage of CD3^+^ T cells), and neutrophils (right panel; as a percentage of total leukocytes) after 2 weeks of control or GC treatment in wild type, *Gzmb^Cre^ Nr3c1^fl/fl^*and *Lyz2^Cre^ Nr3c1^fl/fl^* mice (n=5 per group). (**B**) Growth profiles of melanomas implanted in wild type (left), *Gzmb^Cre^ Nr3c1^fl/fl^* (middle) and *Lyz2^Cre^ Nr3c1^fl/fl^*(right) mice following control or GC treatment (n=5 per group). Data are expressed as mean ± SEM; one-way ANOVA (A) or two-way ANOVA (B). *, *P* < 0.05; **, *P* < 0.01; ****, *P* < 0.0001; ns, not significant. Arrow indicates start of treatment.

**Supplementary Figure 8.**
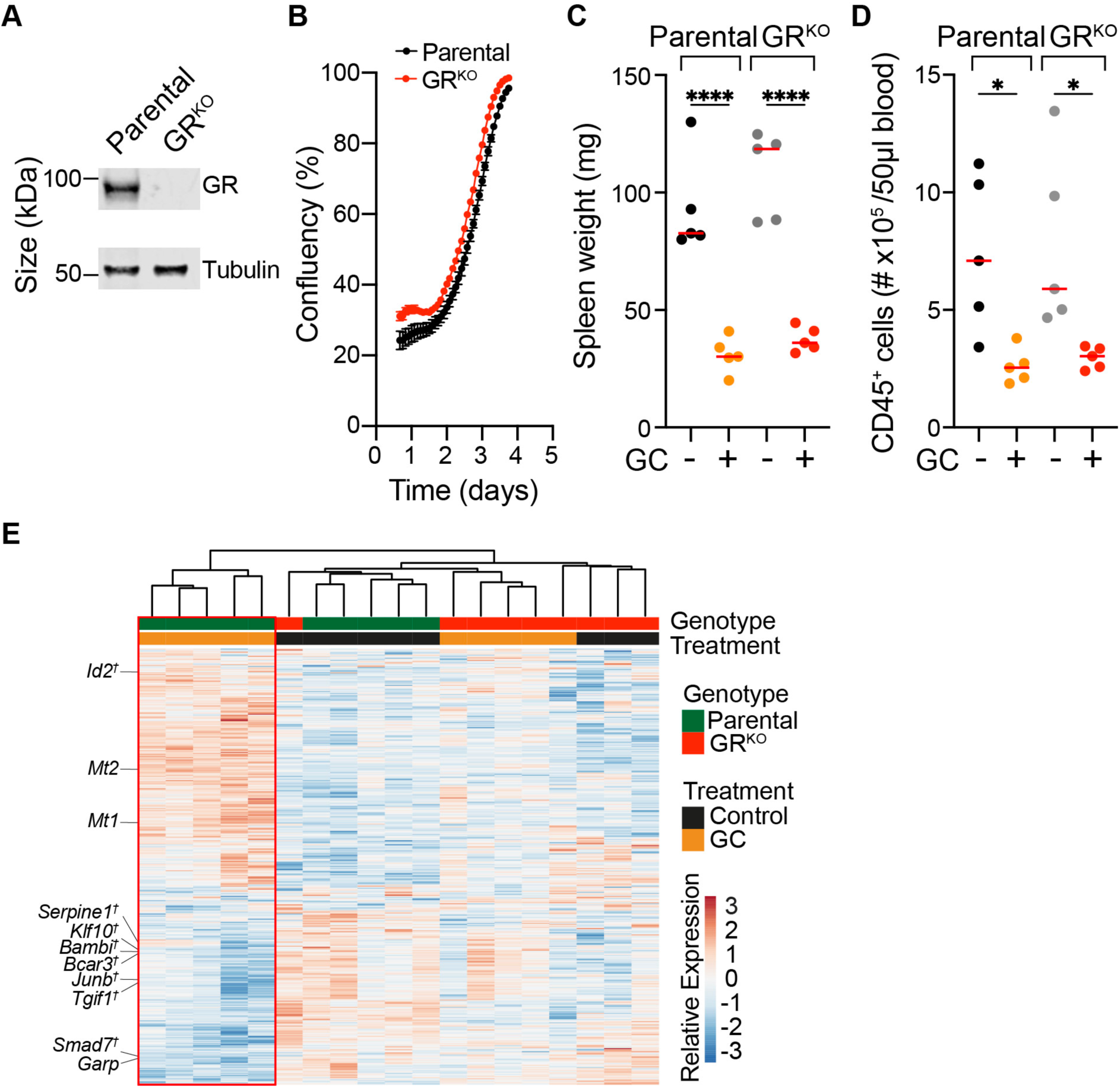
Topical GC treatment induces comparable systemic leukocyte reduction in parental and GR^KO^ tumor-bearing mice. (**A**) Western blot of GR in parental and GR^KO^ 20967 melanoma cells. (**B**) *In vitro* growth of parental and GR^KO^ 20967 melanoma cells. (**C**) Spleen weight 5 days post treatment start with control or GC (n=5 per group). (**D**) Peripheral blood CD45^+^ cell count at day 14 on GC or control treatment (n=5 per group). (**E**) Unsupervised clustering of differentially expressed genes (DEGs) in parental or GR^KO^ tumors with or without GC treatment following RNA sequencing *in vivo* (n=4-5 mice per group). *Mt1* and *Mt2* (GC-inducible), *Garp*, and members of Hallmark TGF-β signaling gene set^†^ indicated. Data are expressed as mean ± SEM; one-way ANOVA (C,D). *, *P* < 0.05; ****, *P* < 0.0001; ns, not significant.

**Supplementary Figure 9.**
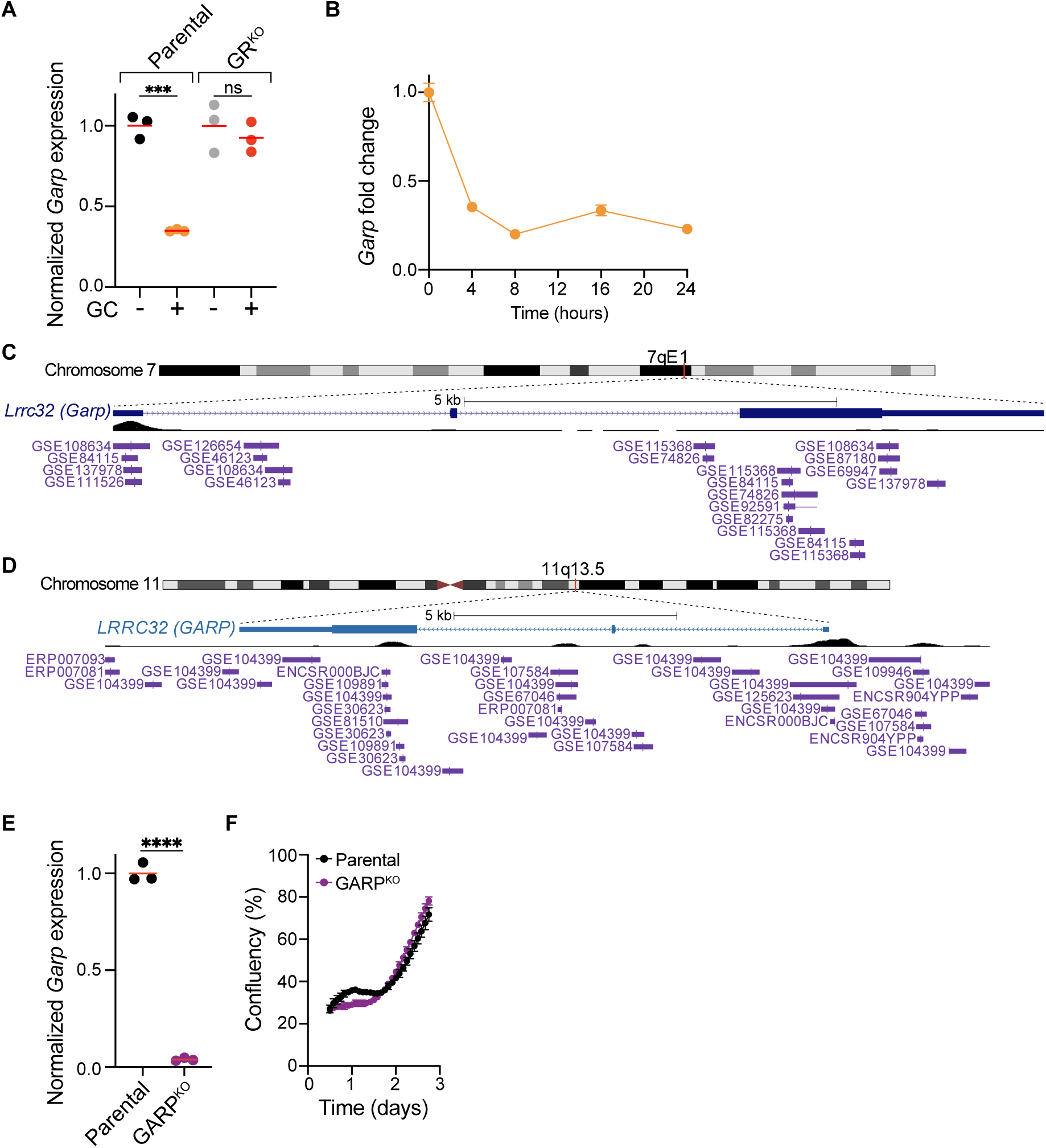
GARP is a direct transcriptional target of GCs. (**A**) GARP mRNA expression by qPCR of parental and GR^KO^ melanoma cells 24-hours post GC treatment *in vitro*. Data are expressed as normalized to *Hprt* and the untreated cell population. (**B**) Time course analysis of GARP mRNA expression by qPCR over 24 hours in GC-treated parental 20967 melanoma cells. Data are expressed as normalized to *Hprt* and baseline GARP levels. (**C, D**) Ch-IP sequencing data showing binding sites of GR (purple) in GARP (Lrrc32) gene in mice (C) and humans (D) in indicated published studies (purple labels). (**E**) GARP expression in parental and GARP^KO^ cells by PCR. (**F**) *In vitro* growth of parental and GARP^KO^ 20967 melanoma cells. Data are expressed as mean ± SEM; one-way ANOVA (A) and unpaired t-test (E). ***, *P* < 0.001; ****, *P* < 0.0001; ns, not significant.

**Supplementary Figure 10.**
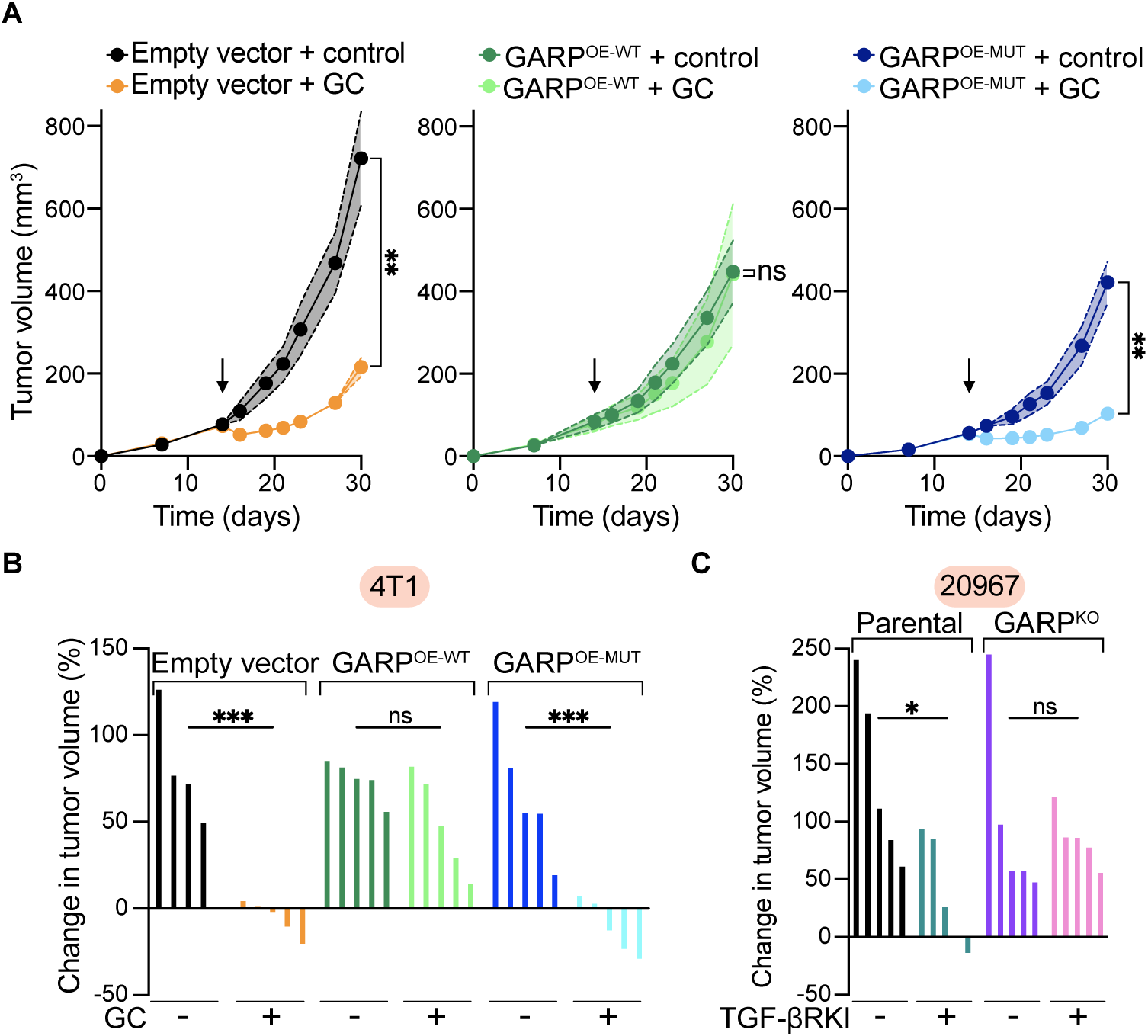
The TGF-β activating function of GARP is required for GC-induced tumor control. (**A**) Growth profiles of empty vector (left), GARP^OE-WT^ (middle) and GARP^OE-MUT^ (right) melanomas following control or GC treatment (n=5 per group). Arrow represents start of treatment. (**B**) Waterfall plots showing percentage change in tumor volume on day 5 of treatment of control and GC-treated empty vector, GARP^OE-WT^ and GARP^OE-MUT^ 4T1 breast cancer tumors (n=5 per group). (**C**) Waterfall plots showing percentage change in tumor volume on day 5 of treatment of control and TGF-βRKI-treated parental or GARP^KO^ melanomas (n=5 per group). Data are expressed as mean ± SEM; two-way ANOVA (A) and one-way ANOVA (B, C). *, P<0.05; ***, *P* < 0.001; ns, not significant.

**Supplementary Figure 11.**
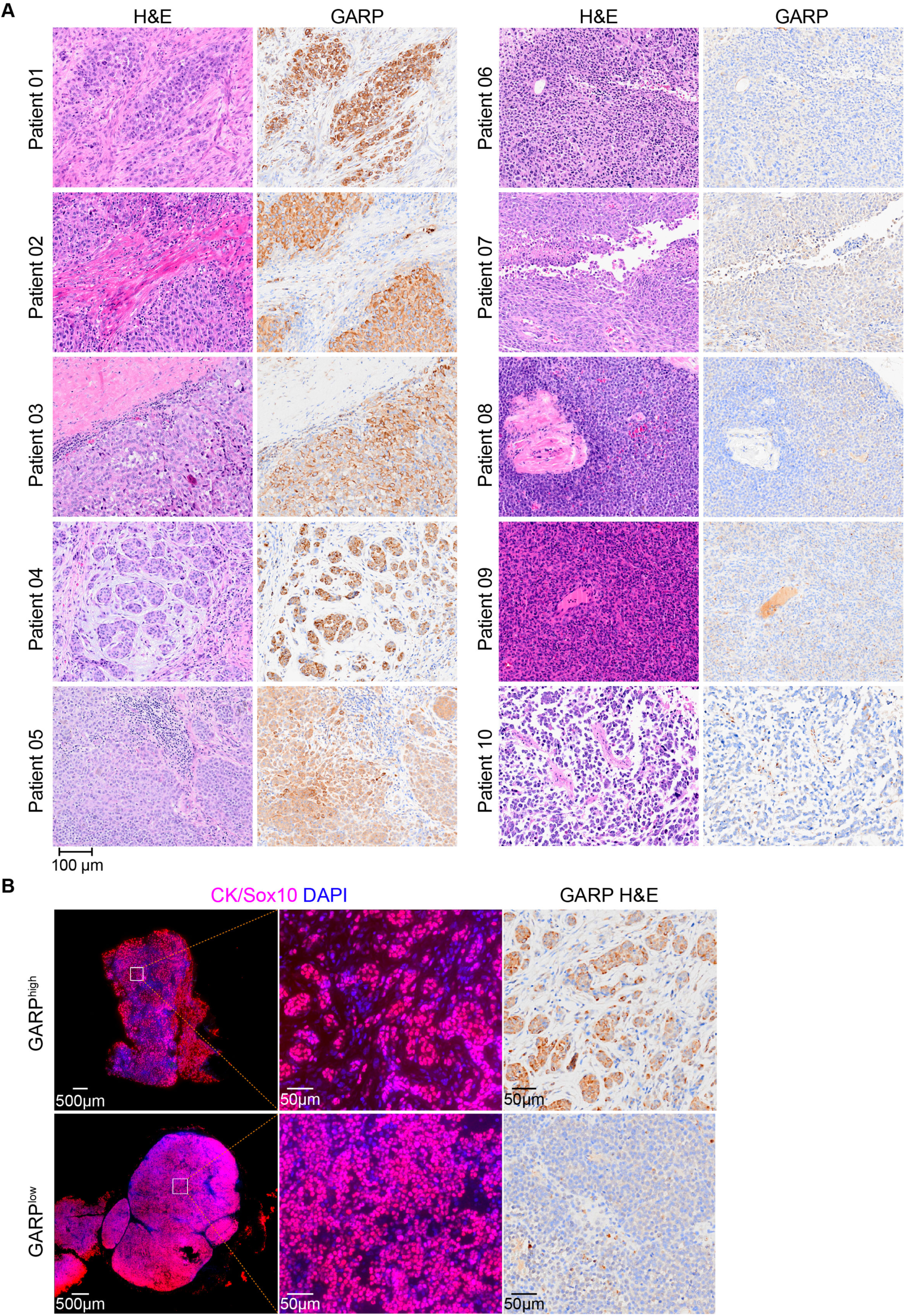
Cancer cell-intrinsic GARP expression is markedly heterogeneous across melanoma patients. (**A**) Representative immunohistochemistry images of 10 melanoma patients (patients 1-5 demonstrating high GARP levels; patients 6-10 demonstrating low GARP levels; Patients 1&2 and 5&6 shown in Fig. 7A). (**B**) Staining of CK/Sox10, DAPI and GARP on serial sections of patient melanoma biopsies.

**Supplementary Figure 12.**
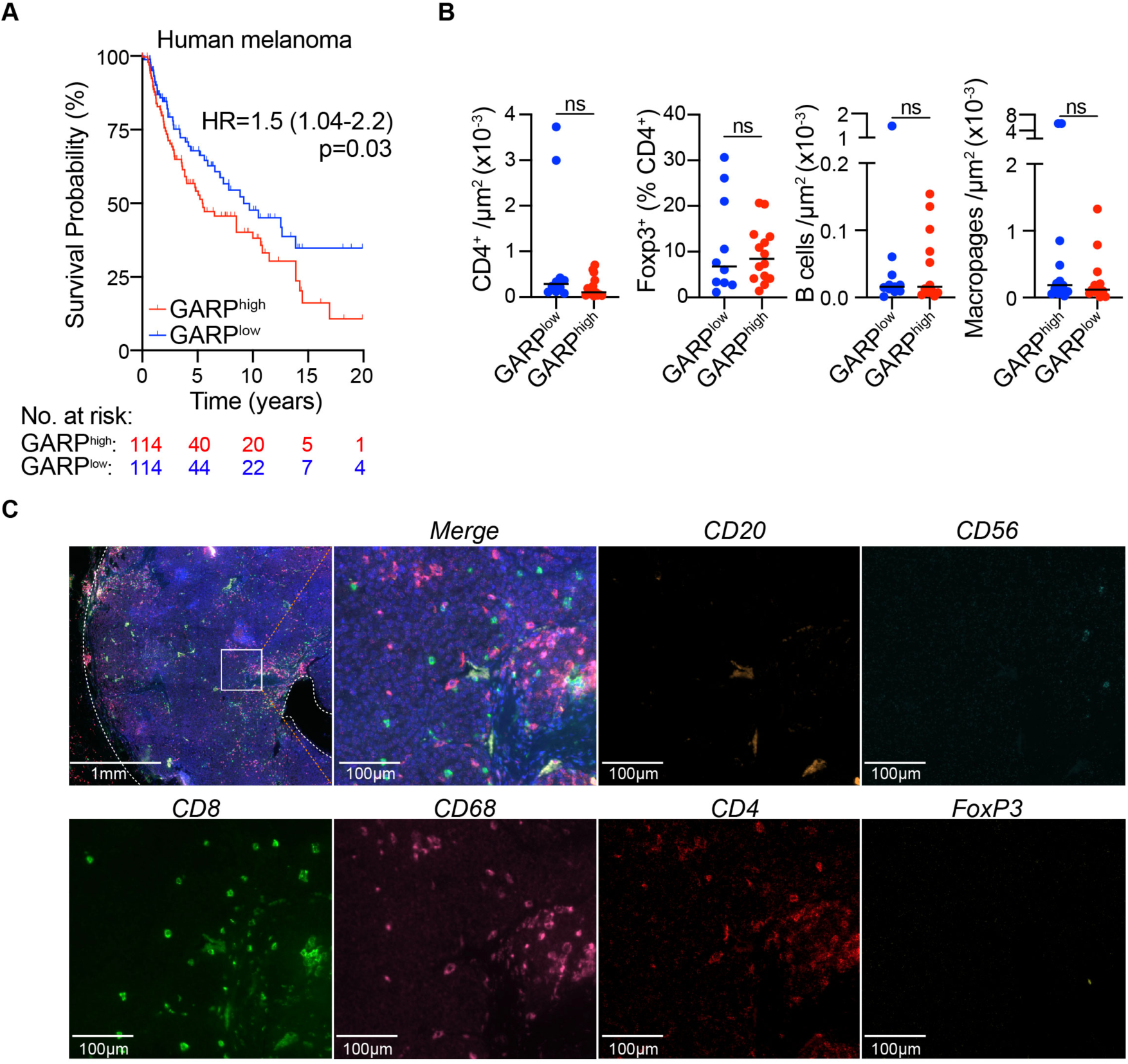
GARP^high^ tumors have poorer prognosis but broadly comparable immune infiltrate composition. (**A**) Kaplan-Meier survival plot of TCGA melanoma patients (n=458) stratified by upper and lower quartile of GARP expression. (**B**) Analysis of in-house melanoma samples from Figure 6B by multiparametric immunofluorescence. Prevalence of indicated populations shown. (**C**) Representative images from human multiplex immunofluorescence analysis of in-house melanoma cohort (Figure 6B). Hazard ratio (95% confidence interval), log-rank (Mantel-Cox) test (A), unpaired t-test (B). ns; non-significant.

**Supplementary Table 1 (separate file).**

Supplementary Table 1. Overlapping DEGs following GC-treatment *in vivo* and *in vitro*, leading to development of GC-stimulated gene signature (related to Fig. 4).

**Supplementary Table 2.**

Supplementary Table 2. Human GC-stimulated gene signature, developed from murine sequencing data (related to Supplementary Table 1).

**Table S3. (separate file)**

List of DEGs in GC-treated parental and GR^KO^ 20967 melanoma cells *in vitro* related to Fig. 4.

**Supplementary Table 4. (separate file)**

GSEA results for GC-treated parental and GR^KO^ tumors *in vivo* related to Fig. 4.

**Supplementary Table 5. (separate file)**

Cluster biomarkers used for cell classification (Fig. 3).

**Supplementary Table 6. (separate file)**

GSEA results for control- or GC-treated 20967 melanomas for the CD8, neutrophil and monocyte clusters related to Fig. 3.

## Notes

**Conflicts of Interest:** SZ received research grants from Ono Pharmaceutical and Nxera Pharma, and honoraria from Ribonexus, Medibiofarma, iTEOS Therapeutics and Owkin, all unrelated to this work. CEMG received research grants from Almirall and Boehringer Ingelheim and received honoraria from Abbvie, Almirall, Boehringer Ingelheim, Boots Ltd, Bristol Meyers Squibb, Evelo Bioscience, GSK, Inmagene, Janssen, Lilly, Novartis, and Ono Pharmaceuticals, all unrelated to this work. The other authors declare they have no competing interests.

### Competing Interest Statement

SZ received research grants from Ono Pharmaceutical and Nxera Pharma, and honoraria from Ribonexus, Medibiofarma, iTEOS Therapeutics and Owkin, all unrelated to this work. CEMG received research grants from Almirall and Boehringer Ingelheim and received honoraria from Abbvie, Almirall, Boehringer Ingelheim, Boots Ltd, Bristol Meyers Squibb, Evelo Bioscience, GSK, Inmagene, Janssen, Lilly, Novartis, and Ono Pharmaceuticals, all unrelated to this work. The other authors declare they have no competing interests.

